# Calpains Orchestrate Secretion of Annexin-containing Microvesicles during Membrane Repair

**DOI:** 10.1101/2024.09.05.611512

**Authors:** Justin Krish Williams, Jordan Matthew Ngo, Abinayaa Murugupandiyan, Dorothy E. Croall, H Criss Hartzell, Randy Schekman

## Abstract

Microvesicles (MVs) are membrane-enclosed, plasma membrane-derived particles released by cells from all branches of life. MVs have utility as disease biomarkers and may participate in intercellular communication; however, physiological processes that induce their secretion are not known. Here, we isolate and characterize annexin-containing MVs and show that these vesicles are secreted in response to the calcium influx caused by membrane damage. The annexins in these vesicles are cleaved by calpains. After plasma membrane injury, cytoplasmic calcium-bound annexins are rapidly recruited to the plasma membrane and form a scab-like structure at the lesion. In a second phase, recruited annexins are cleaved by calpains-1/2, disabling membrane scabbing. Cleavage promotes annexin secretion within MVs. Our data supports a new model of plasma membrane repair, where calpains relax annexin-membrane aggregates in the lesion repair scab, allowing secretion of damaged membrane and annexins as MVs. We anticipate that cells experiencing plasma membrane damage, including muscle and metastatic cancer cells, secrete these MVs at elevated levels.

## Introduction

Extracellular vesicles (EVs) are membrane-enclosed compartments secreted by cells from all three branches of life. EVs are divided into two subtypes: Microvesicles (MVs) that bud directly from the plasma membrane, and exosomes that form intracellularly. EVs have utility as biomarkers, as they contain protein and RNA specific to their origin cell. EVs may also function in intracellular communication (1). While significant attention has been devoted to exosomes, the mechanisms and physiological circumstances driving MV secretion are poorly understood.

EVs are enriched in annexins (2), which bind to phospholipids in the presence of calcium. Annexins have roles in plasma membrane repair. Knockdowns of several individual annexins cause defects in membrane resealing in cultured cells (3, 4). Muscle cells experience high rates of membrane damage *in vivo* due to repeated contractions and are dependent on the membrane repair machinery for survival. For example, 20% of muscle fibers in rat triceps are disrupted after eccentric exercise (5). Consistent with annexins playing a role in muscle repair, annexin mutations cause muscular dystrophy in mice (6, 7). Upon plasma membrane damage, extracellular calcium flows into the cytoplasm. In the presence of micromolar calcium, annexins bind to membranes and may plug lesion sites by tethering membranes (8). These membranes may come from intracellular organelles or from reticulation of the plasma membrane (9). After annexin recruitment, annexin^+^ MVs are shed from the lesion (10). Recently, Jeppesen et al. separated annexin-rich MVs from exosomes using density gradient centrifugation (2). The contents, mechanism of their shedding, and roles of annexin-rich MVs in membrane repair are not understood.

Calpains are cytosolic, calcium-dependent cysteine proteases that, like annexins, have roles in membrane repair. Calpain-1/μ-calpain and calpain-2/m-calpain were the first characterized calpains and are highly abundant across tissues. Calpain-1 and calpain-2 have separate large subunits but share a small subunit. Calpain-3 – calpain-14 are less abundant and may be restricted to specific tissues (11). Like many annexin knockouts, loss of calpain-1 and calpain-2 leads to impaired membrane resealing in cell culture (12) and severe muscular dystrophy in mice (13). Mutations in the muscle specific calpain-3 produce limb-girdle muscular dystrophy type R1 (8). As with annexins, calpains are activated by micromolar concentrations of calcium *in vitro*. Because calpain targets include membrane-cytoskeleton anchors, it has been speculated that calpain-1/2 locally detach the plasma membrane from the cytoskeleton around the lesion site (14). In other systems, calpains also appear to be necessary for MV formation. Platelets release MVs called microparticles during activation and clotting. Microparticle formation in platelets requires sustained elevation of cytosolic calcium and is blocked by calpain inhibitors (15). It is unknown if calpains cleave other proteins and facilitate repair and MV formation independently of membrane-cytoskeleton detachment.

Membrane repair appears to be important for cancer cell migration and metastasis (16). To characterize membrane repair and mechanisms of MV secretion in cancer cells, we isolated and identified proteins within annexin^+^ MVs produced by cells grown in culture. After finding several calcium-activated proteins implicated in membrane repair in MVs, we measured annexin secretion in MVs after membrane damage. Addition of sublytic levels of the pore forming toxin, streptolysin O, induced ∼20-40-fold increase in vesicular annexin A2 secretion. We found that secreted annexin A1 and annexin A2 are cleaved by calpain-1/2, and that annexins in general are primary targets for calpains. Given that both annexins and calpains are required for membrane repair, we sought to understand the interplay of these proteins during membrane repair. Using an intracellular annexin A2 cleavage reporter, we found that annexin A2 was cleaved after membrane recruitment and wound scabbing, but before secretion. We show that cleaved annexin A2 is deficient in membrane and effector protein binding, and cleavage dissociates annexin A2-linked membranes. Mutant annexin A2 that cannot be cleaved by calpains was recruited normally after membrane damage but was secreted at lower levels in MVs compared to wildtype annexin A2. Our results link membrane repair to MV secretion and establish a chronology of annexin and calpain function during membrane repair.

## Results

### Annexins and other calcium-responsive proteins are secreted within microvesicles

To characterize microvesicles (MVs) secreted in culture, we utilized serial centrifugation followed by equilibrium density gradient centrifugation. This established method separates EVs based on their buoyant density (17). Following a 1k x g centrifugation to remove cells and further centrifugation at 10k x g to remove large EVs, small EVs were sedimented from conditioned medium at 100k x g. Sedimented particles were resuspended and placed at the bottom of an iodixanol gradient and centrifuged to separate HCT116-derived small EVs into a low-density, annexin^+^ population and a high-density, CD63^+^ population (Fig. 1a). The CD63^+^ population is thought to represent exosomes, and the annexin A2^+^ population is thought to represent a subpopulation of MVs (2). Vesicles centrifuged from conditioned medium were immunoprecipitated with CD63 antibody (Figure 1b). Known exosome markers, CD9 and alix, coprecipitated with CD63^+^ vesicles, but annexin A1 did not. We assessed whether annexins were localized to the lumen or the extracellular face of MVs by proteinase K treatment. We found that annexin A2 mostly localized to the lumen of MVs (Figure 1c). Luminal flotillin-2 and annexin A2 sedimented with EVs even in the presence of the membrane impermeable calcium chelator, EGTA, and both were degraded by proteinase k only in the presence of triton X-100.

**1.**
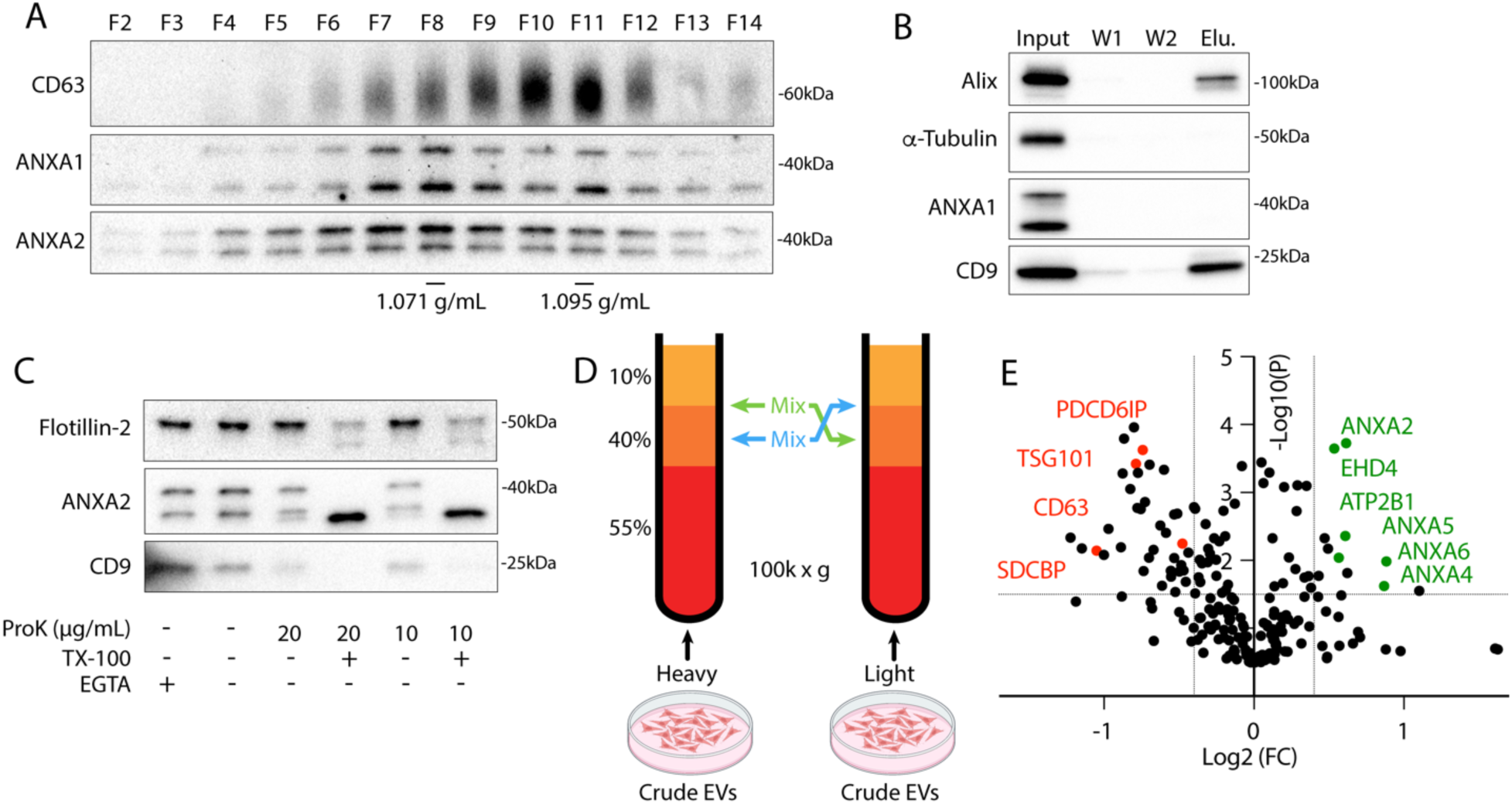
Annexin-containing extracellular vesicles are distinct from exosomes. (A) Immunoblots show distribution of EV markers across an iodixanol gradient of the conditioned medium 100k × g pellet fraction. Samples were taken from low (F2-Fraction #2) to high density (F14-Fraction #14). (B) Immunoblots show depletion of annexins from exosomes after immunoprecipitation with anti-CD63 beads from the conditioned medium 100k × g pellet fraction (W1-wash #1, W2-wash #2). (C) Immunoblots show degradation of EV markers in the conditioned medium 100k × g pellet fraction after treatment with indicated combinations of proteinase K (ProK) and 0.1% Triton X-100 (TX-100). For EGTA treatments, 5 mM EGTA was introduced into the medium prior to centrifuging the conditioned medium at 100k × g. (D) Schematic illustrating the separation of EVs for quantitative proteomics using sucrose step gradients is shown. (E) Volcano plot shows enriched proteins in the high buoyant density fractions (red) relative to the low buoyant density fractions (green). P-values were calculated using a t-test.

Using stable isotope labeling by amino acids in cell culture (SILAC), we identified proteins enriched in the low-density fraction relative to the high-density fraction. EVs secreted by HEK293T cells grown with heavy arg/lys or light arg/lys amino acids were collected from conditioned medium, centrifuged at 100k x g, and pellet material was resuspended and placed beneath a sucrose step gradient (Figure 1d). After separation by high-speed centrifugation (Fig. S1a), heavy amino acid labeled, low buoyant density EVs, collected from the 10-40% sucrose interface, were mixed with light amino acid labeled, high buoyant EVs, collected from the middle of the 40% sucrose fraction (and vice versa) prior to mass spectrometry analysis. As expected, conventional exosome proteins (CD63, ESCRTs, syntenin-1, etc.) were enriched in the high buoyant density fraction (Figure 1e, Table S1). Conversely, in the low buoyant density fractions Ca^2+^-responsive genes including annexins, EHD proteins (EHD1, EHD4), plasma membrane snare machinery (SNAP23, STXBP3), and the Ca^2+^-transporting ATPase (ATP2B1) were enriched. Annexin, EHD (18), and snare (19) proteins are all implicated in plasma membrane repair. As expected for plasma membrane derived vesicles, we also detected several plasma membrane proteins including CD276 and CD59.

### Membrane damage stimulates secretion of annexin A2^+^ microvesicles from the repair site

We showed previously that in addition to exosomes, secretion of this annexin A2^+^ EV population is stimulated by the Ca^2+^ ionophore, ionomycin (20). In addition, we have previously visualized secretion of annexin ^+^ MVs from muscle cells during Ca^2+^-dependent membrane repair (10). Given these previous data, and our proteomics results, we speculated that secretion of these vesicles might be stimulated by membrane damage. First, we visualized (Figure 2a, Video 1) and quantified (Figure 5h) FM1-43 staining of HCT116 cells after laser ablation wounding. FM1-43 is membrane impermeable but can stain intracellular membranes brightly if the plasma membrane is disrupted. Within 1 min of ablation, a repair scab of brightly stained membranes formed at the ablation site. For several min after ablation, intracellular membranes slowly stained, suggesting the lesion site was plugged but not fully sealed. As dye influx slowed, vesicles were shed from the damaged cell. These vesicles were intensely stained at a level comparable to the repair site, possibly indicating they were shed directly from the repair site. In addition, as vesicles were shed over the next 5 min, the repair scab slowly lost staining intensity.

**2.**
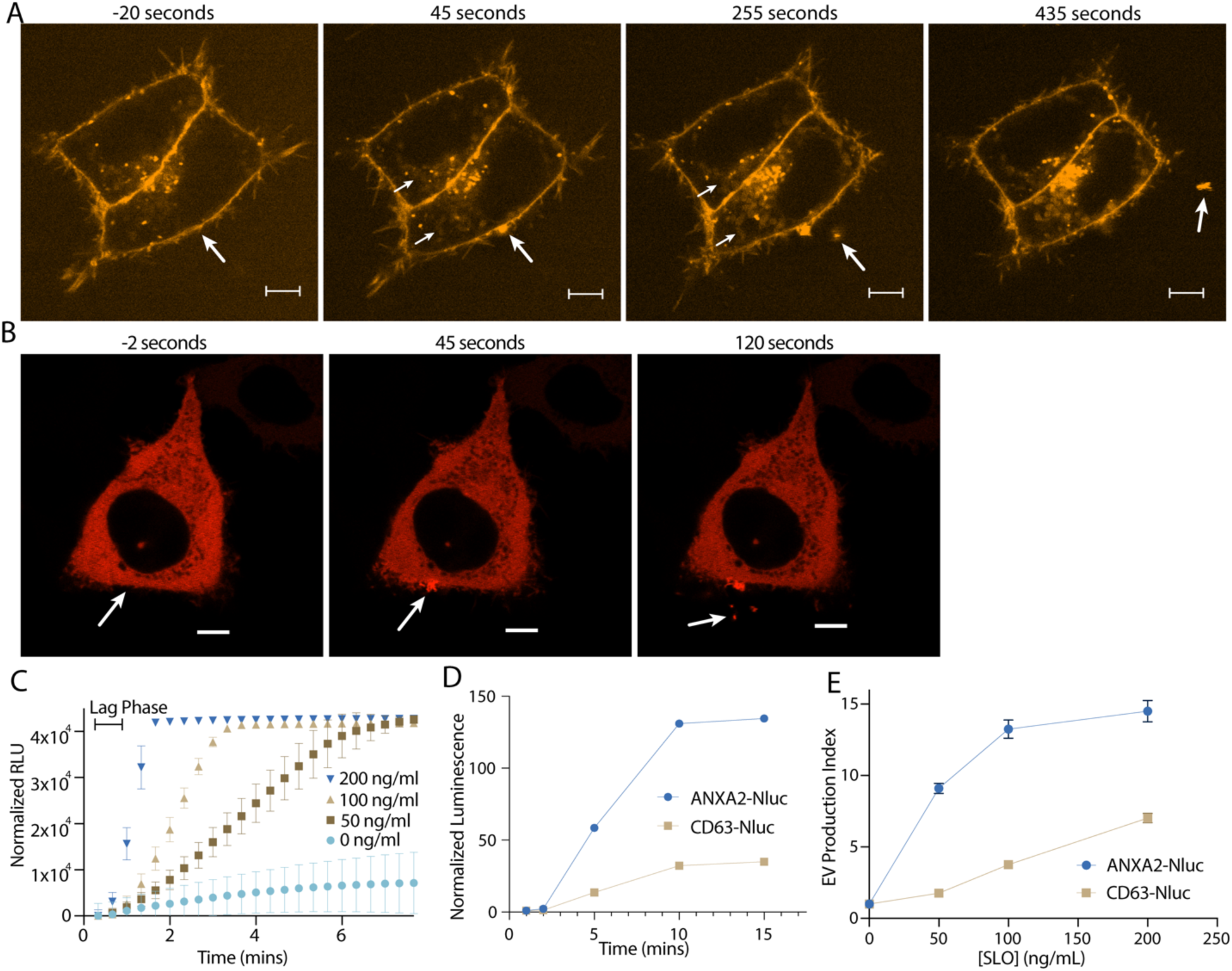
Annexin-containing extracellular vesicles are shed from the repair scab after plasma membrane damage. (A) Representative confocal micrographs of FM1-43 infiltration are shown. Image times are relative to the first image taken post ablation. Large arrows in pane I and II indicate the site of ablation. Small arrows in pane II and III indicate the staining of intracellular compartments in an ablated (lower arrows) and a non-ablated cell (upper arrows). Large arrows in pane III and IV indicate extracellular vesicles. Scale Bars: 5 μm. (B) Representative confocal micrographs of ANXA2-mScarlet recruitment after ablation are shown. Image times are relative to the first image taken post ablation. White arrows in pane I and II indicate the site of ablation. Arrow in pane III indicates an extracellular vesicle. Scale Bars: 5 μm. (C) Sytox Green staining over time in the absence of extracellular Ca^2+^ after cells pretreated with the indicated concentration of SLO were rapidly heated from 4°C to 37°C. (D) Membrane-protected luminescence in medium fractions at indicated time points after ANXA2-Nluc or CD63-Nluc cells pretreated with 200 ng/μl SLO were rapidly heated from 4°C to 37°C is shown. (E) EV production index from ANXA2-Nluc or CD63-Nluc cells treated with increasing concentrations of SLO is shown.

Next, we visualized annexin dynamics after laser ablation (Figure 2b). We focused on annexin A2, as it is the most abundant annexin in many cell types. Less than 1 min after ablation, annexin A2 was rapidly recruited to the membrane around the lesion site. For the next 5-10 min after recruitment, annexin A2^+^ vesicles were shed from the ablation site. As these vesicles were shed, the membrane-recruited annexin A2 dissipated. The morphology of the repair scab varied (Fig. S2a). Occasionally, annexin A2^+^ filopodia-like structures emerged rapidly after ablation, before shedding from the damage site. We also observed size variation in annexin A2^+^ EVs, ranging from 1 μM (Fig. S2b) to 100-200 nM (Figure 2b). In these larger vesicles, we could distinguish complete recruitment of annexin A2 to the membrane, suggesting that the Ca^2+^ concentration in these vesicles was significantly higher than the intracellular concentration.

To quantitatively access vesicle shedding during membrane repair, we used the pore forming toxin, streptolysin-O (SLO), to damage the plasma membrane. We began to observe increased cell death at 800 ng/ml SLO as assessed by SYTOX Green staining after treatment (Fig. S2c). Thus, we considered treatments of 200 ng/ml SLO and less, sublytic. We synchronized pore formation by pre-incubating cells at 4°C with SLO. At this temperature, SLO binds to the plasma membrane but does not open into a pore (21). Cells were washed to remove excess SLO, after which pores open synchronously as the temperature was raised to 37°C.

To assess the dynamics of pore formation using this system, we measured uptake of SYTOX Green into HCT116 cells in Ca^2+^-free medium. Under these conditions, SLO pores remained open and were not repaired (21). SLO pores opened within 40 sec to 1 min, as measured by the length of the lag phase in SYTOX Green uptake (Figure 2c). Next, we applied SLO to cells expressing a low level of annexin A2-nanoluciferase (Nluc). Using an assay described previously that measures luminal luciferase activity (20), we quantified secretion of membrane enclosed annexin A2-Nluc over time (Figure 2d, S2d). Neither annexin A2-Nluc nor CD63-Nluc were secreted within the first two min of SLO treatment. By 5 min, both annexin A2-Nluc and CD63-Nluc secretion peaked, and by 10 min secretion had ceased. Accounting for the 1 min it took for SLO pores to open, vesicle secretion initiated 1 min after pore formation and peaked after about 4 min. Next, we measured annexin^+^ MV secretion at increasing concentrations of SLO (Figure 2e). Annexin A2-Nluc reached maximal secretion with 100-200 ng/ml SLO whereas CD63-Nluc, a marker of exosomes, did not reach maximal secretion within the tested concentration range. Annexin A2^+^ MV secretion was stimulated at lower levels of damage, whereas late endosome exocytosis is required at higher levels of damage.

Membrane repair and repair cap shedding involves the phospholipid scramblases, TMEM16F or TMEM16E (10, 22). We speculated that annexin^+^ EVs may have high levels of anionic phospholipids, such as phosphatidylserine, on the extracellular leaflet if they are derived by budding from the plasma membrane repair site. We found that annexin A1^+^ and A2^+^ EVs were more efficiently captured by an immobilized form of the phosphatidylserine-binding protein annexin A5 compared to CD63^+^ and Alix^+^ exosomes (Fig. S2e). Thus, annexin^+^ EVs are enriched in phosphatidylserine and/or phosphatidylethanolamine in their outer membrane leaflet.

Membrane repair also requires Endosomal Sorting Complexes Required for Transport (ESCRT) machinery (23). Although some reports suggest ESCRTs and annexins work in tandem during membrane repair (4), others have reported independent activity (23). ESCRT-mediated vesicle fission depends upon the ATPase activity of Vps4. Inducing short term expression of dominant negative VSP4a diminished vesicular annexin A2-Nluc signal compared to the uninduced control (Fig. S2f). We conclude that annexin^+^ MV shedding during membrane repair at least partially depends on the ESCRT pathway.

### Annexins within microvesicles have altered apparent molecular weight

Next, we examined the role of extracellular Ca^2+^ in secretion of these annexin^+^ vesicles. Significant levels of annexin proteins were detected in buoyant fractions of a sucrose density gradient, but only when Ca^2+^ was present in the medium (Figure 3a). We noted that annexin A1 and A2 within MVs migrated at a lower apparent molecular weight. No such annexin mobility shifts were detected in cell lysates even after SLO treatment (Figure 3b). We confirmed that annexin proteins were secreted in a distinct vesicle fraction from exosomes after SLO treatment. As expected annexin A1, A2, and A6 were present in a low-density fraction, relative to high-density, CD63^+^ exosomes (Figure 3c). Annexin A6 also appeared to shift in molecular weight, although the apparent molecular weight change was much larger than for annexin A1 or A2. Because muscle cells experience some of the highest rates of membrane damage, we examined this effect using C2C12 cells. As with tumor cells, myotubes, differentiated from C2C12 cells, secreted EVs with apparently processed annexins in response to SLO treatment (Fig. S3a).

**3.**
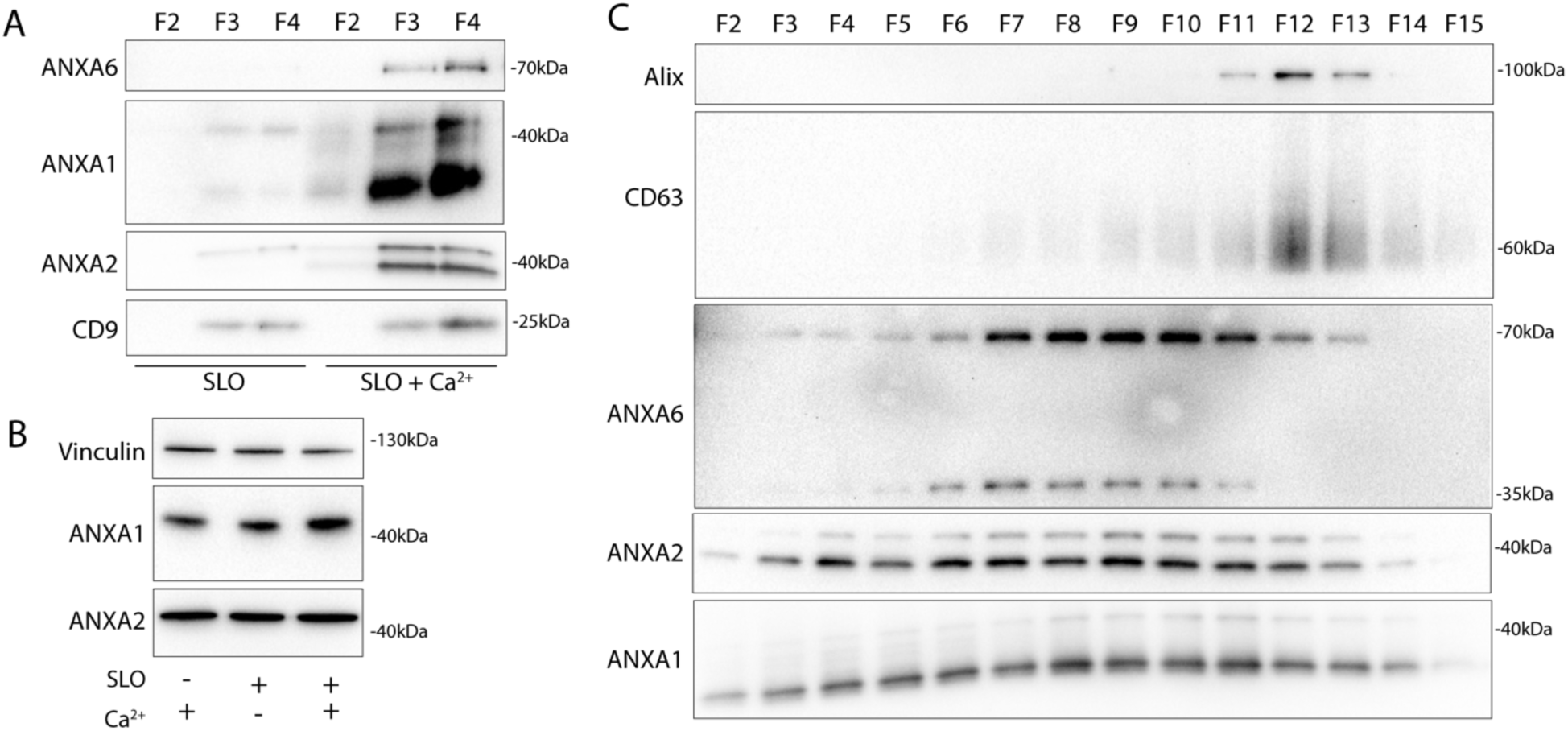
Annexins within extracellular vesicles are shifted in apparent molecular weight. (A) Immunoblots of gradient fractions show enrichment of indicated EV markers after 200 ng/mL SLO treatment, with or without 1 mM Ca^2+^ in the media. F2, F3, and F4 refer to buoyant fractions of a sucrose step gradient of the conditioned medium 100k × g pellet fraction. (B) Immunoblot analysis of cell lysates after treatment with indicated combinations of 1 mM Ca^2+^ and 200 ng/mL SLO is shown. (C) Immunoblots show separation of indicated EV markers after 200 ng/mL SLO treatment. F2-F15 refer to fractions #2-15 of an iodixanol gradient of the conditioned medium 100k × g pellet fraction, moving from low to high density.

We speculated that SLO pores may be present on annexin^+^ vesicles derived from the secreted plasma membrane repair scab. Indeed, others reported such pores on vesicles secreted from SLO-treated cells (24). In our experiments, SLO was secreted predominantly on larger vesicles that sedimented at 10k x g (Fig. S3b). We reasoned that such perforated vesicles would be accessible to membrane impermeable compounds. As expected, a larger fraction of annexin A2-Nluc secreted in larger vesicles was accessible to a membrane impermeable Nluc inhibitor compared to annexin A2-Nluc secreted in smaller vesicles (Fig. S3c). We conclude that cells secrete both perforated and sealed EVs in response to membrane damage.

### Calpain-1/2 proteases cleave annexins during microvesicle shedding

We investigated the basis of the apparent molecular weight change of the annexins in MVs. Annexins are substrates for phosphorylation by kinases including Src or PKC (25), and for proteolysis by enzymes including metalloproteases (26), plasmin, cathepsins, and calpains (27). We tested the role of Ca^2+^ in the apparent molecular weight change for annexins and found that addition of 1 mM in a lysate of HCT116 cells resulted in a rapid conversion of annexin A2 (Figure 4a).

**4.**
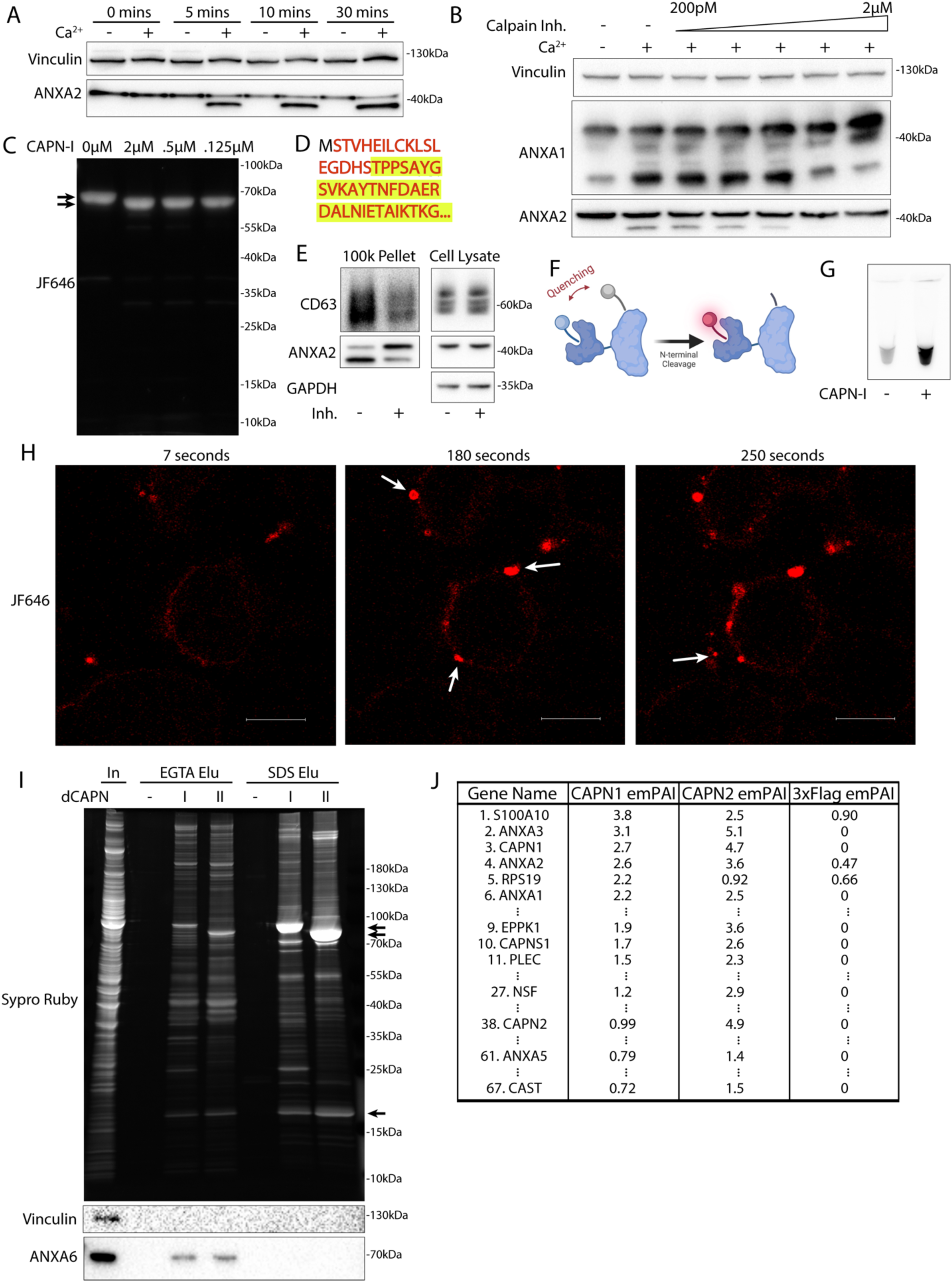
Calpain-1/2 cleave annexins which are then shed in microvesicles. (A) Immunoblot analysis of cytosol fractions after incubation with or without 1 mM Ca^2+^. (B) Immunoblot analysis of cytosol fractions after incubation with or without 1 mM Ca^2+^, with a range of concentrations of calpastatin domain I inhibitor (Calpain Inh.). (C) In gel fluorescence of JF646-labeled, recombinant annexin A2-Halo incubated with indicated concentration of purified, porcine calpain-1. Arrows indicate uncleaved and cleaved product. (D) Mapping of tryptic peptides to the first 50 amino acids of annexin A2. Mass spectrometry analysis of recombinant annexin A2 (red text) is compared to tryptic digest-mass spectrometry analysis of recombinant annexin A2 treated with purified, porcine calpain-1 (yellow highlight). (E) Immunoblot analysis of cell lysate and conditioned medium 100k × g pellet fraction after treating cells with 200 ng/ml SLO in the presence or absence of 10 μM calpain inhibitor, ALLN. (F) Schematic illustrating the recombinant annexin A2-Halo reporter, labeled with a 5WS maleimide quencher on the N-terminus and a JF646 halo ligand on the C-terminus. (G) JF646 fluorescence of an *in vitro* reaction containing 5 μM self-quenched annexin A2 with or without 0.5 μM porcine calpain-1. (H) Representative confocal micrographs of dequenched annexin A2-Halo-JF646 fluorescence in cells. Times are relative to the first image taken post addition of 1 mM Ca^2+^. White arrows in pane II indicate puncta on the cell periphery. Arrow in pane III indicates an extracellular vesicle. Scale Bars: 10 μm. (I) Total protein (Sypro Ruby staining) and immunoblot analysis of substate trapping experiment, using 3xFlag (−), 3x-Flag C115S calpain-1 (dCAPN-I), or 3x-Flag C105S calpain-2 (dCAPN-II) as bait is shown. Arrows indicate calpain proteins. (J) Table listing proteins identified in EGTA elutions from 3xFlag, 3x-Flag C115S calpain-1 (dCAPN-I), or 3x-Flag C105S calpain-2 (dCAPN-II) capture experiments. Proteins are listed by Exponentially Modified Protein Abundance Index (emPAI) in the dCAPN-I elution. Keratin proteins were not included in the list.

Calpains are a well characterized family of Ca^2+^-dependent proteases that are necessary for membrane repair in cells. Calpains 1 & 2 are the highest expressed calpains across tissues and have ∼10 fold higher transcript levels in HCT116 cells compared to other calpains (28). Addition of the highly specific calpain 1 & 2 inhibitor, calpastatin domain I (K_i_ = 15nM), inhibited the apparent molecular weight shift at a concentration near the K_i_ (Figure 4b). The less specific, membrane permeable calpain inhibitor, ALLN (K_i_ = 200nM), also inhibited the apparent molecular weight shift near the K_i_ (Fig. S4a). We purified annexin A2-halotag using a one step, tag-free purification method (Fig. S4b). Annexin A2-halotag gel mobility-shifted in apparent molecular weight from ∼70kDa to ∼67kDa when combined with purified calpain-1 and Ca^2+^ (Figure 4c). The cleavage appeared to be remarkably specific, producing only the 67kDa species, even in incubations at equal concentrations of calpain and annexin A2-halotag (2μM). Purified calpain-1 also cleaved recombinant annexin A6 into 35kDa products (Fig. S4c, S4d) which matched the shift in apparent molecular weight of annexin A6 in MVs (Figure 3c). The consensus sequence that predicts the substrate specificity of calpain-1/2 is not well defined (29). Therefore, we mapped the calpain cleavage site on annexin A2 by mass spectrometry and detected peptides covering the first 50 amino acids of the protein (Figure 4d, Table S2). Mass spectrometry of processed annexin A2 revealed a cleavage that removed the first 18 resulting in a polypeptide starting at residue Thr-19. The conversion of annexin A2 in MVs was blocked in cells treated with ALLN (Figure 4e). We conclude that MV annexins are cleaved by Ca^2+^-dependent calpain-1/2.

Given that both annexins and calpains are important for membrane repair, we sought to understand the interplay between these two proteins. Based on the location of the cleavage site (Ser-18/Thr-19), we designed a reporter for annexin A2 cleavage by calpain (Figure 4f). Purified annexin A2-halotag was labeled with a fluorescent halo ligand and a maleimide fluorescence quencher. As the only accessible cysteine on annexin A2-halotag is Cys8, the cleavable N-terminus could be site-specifically labeled with the quencher. Addition of calpain-1 to the reporter led to a robust increase in fluorescence *in vitro* as the N-terminus containing the quencher was cleaved (Figure 4g). Next, we used this reporter in cells. Cells were treated with SLO and the reporter in the absence of calcium, and the reporter was allowed to diffuse into the cells. After washing the cells, addition of calcium triggered the repair process and annexin A2 cleavage was visualized in the cells as an increase in halotag-JF646 fluorescence (Figure 4h). Within 1-2 min, bright, cleaved annexin A2-halotag puncta formed on distended parts of the cell surface, presumably at repair scabs. After puncta formation, vesicles could be seen ejecting from these puncta. These vesicles were also brightly stained, similar to the repair scab puncta. Thus, annexin A2 cleavage by calpain is highly localized in cells and occurs after annexin A2 recruitment and scabbing at the plasma membrane.

Many proteins are reported substrates for calpains. We wondered if annexin proteins are primary targets for calpains, or if they represent a minor fraction of total. We developed an unbiased, substrate-trapping approach to identify calpain-1/2 substrates (Fig. S4e). Calpains (3xFlag-tagged), inactivated by mutation of the catalytic cysteine to serine, were immobilized on anti-flag beads. Cytosol was incubated with the beads and substrates in the presence of Ca^2+^ bound to the open calpain active site. Addition of EGTA chelated Ca^2+^ to close the active site, which resulted in substrates eluting from the beads. Stable, Ca^2+^-independent interactors were retained. We coexpressed and purified CAPN1[C115S]-3xFlag-6xHis (dCAPN-I) and CAPN2[C105S]-3xFlag-6xHis (dCAPN-II) each in complex with calpain small subunit, CAPNS1(86-268) (Fig. S4f).

Using dCAPN-I and dCAPN-II baits, we captured calpain substrates (EGTA elution) and stable interactors (subsequent SDS elution) from HCT116 cytosol (Figure 4i). Consistent with calpains targeting annexins in the presence of Ca^2+^, EGTA quantitatively eluted annexin A6 from dCAPN-I and dCAPN-II baits. Many known calpain substrates are targeted by both calpain-1 and calpain-2. Consistent with these observations, the gel patterns of the dCAPN-I and dCAPN-II EGTA elution were similar. Next, we analyzed EGTA elutions using high resolution mass spectrometry and ranked proteins by their exponentially modified protein abundance index in the dCAPN-I sample (30) (Figure 4j, Table S3). As expected, the endogenous calpain-1/2 inhibitor, calpastatin, was captured by dCAPN-I and dCAPN-II baits, but not by the 3xFlag alone control. In addition, we recovered plectin (PLEC), a known calpain substrate that links the cytoskeleton to the plasma membrane (31). The significant abundance of PLEC and EPPK1 exclusively in the dCAPN-I and dCAPN-II elutions is consistent with the role suggested for calpains in severing the plasma membrane from the cytoskeleton.

S100A10, annexin A3, annexin A2, and annexin A1 were the top few substrates in the dCAPN-I sample. In addition, we found six other annexins (A4, A5, A6, A7, A11) with lower peptide counts. High levels of all the same annexins were identified in the dCAPN-II sample, but not in the 3xFlag alone control. We also found many novel potential calpain-1/calpain-2 substrates including NSF and AHNAK, the latter of which forms a complex with annexin A2 and S100A10 that may be important for plasma membrane repair (32). Our data suggest that annexins and potentially annexin interactors are a major target for calpain-1/2.

### Calpain cleavage decreases annexin A2’s membrane binding and scabbing activity

We next considered how cleavage changes annexin behavior at the repair site. To investigate this, we designed two annexin A2 mutants. The published structure of calpain-2 with its endogenous inhibitor, calpastatin, suggests two calpastatin prolines are important for recognition (33). Annexin A2 also has two prolines (Pro-27 and Pro-28) directly adjacent to the calpain cleavage site. Mutating this site (P27D/P28D) rendered calpain incapable of cleaving annexin A2 (Figure 5a). We also created a second truncation mutant, annexin A2(18-339) (tr-annexin A2), that reproduced annexin A2 post-cleavage (Figure 5b). For technical reasons, SLO was added to cells at 37°C. This procedure resulted in delayed, asynchronous pore formation and repair.

**5.**
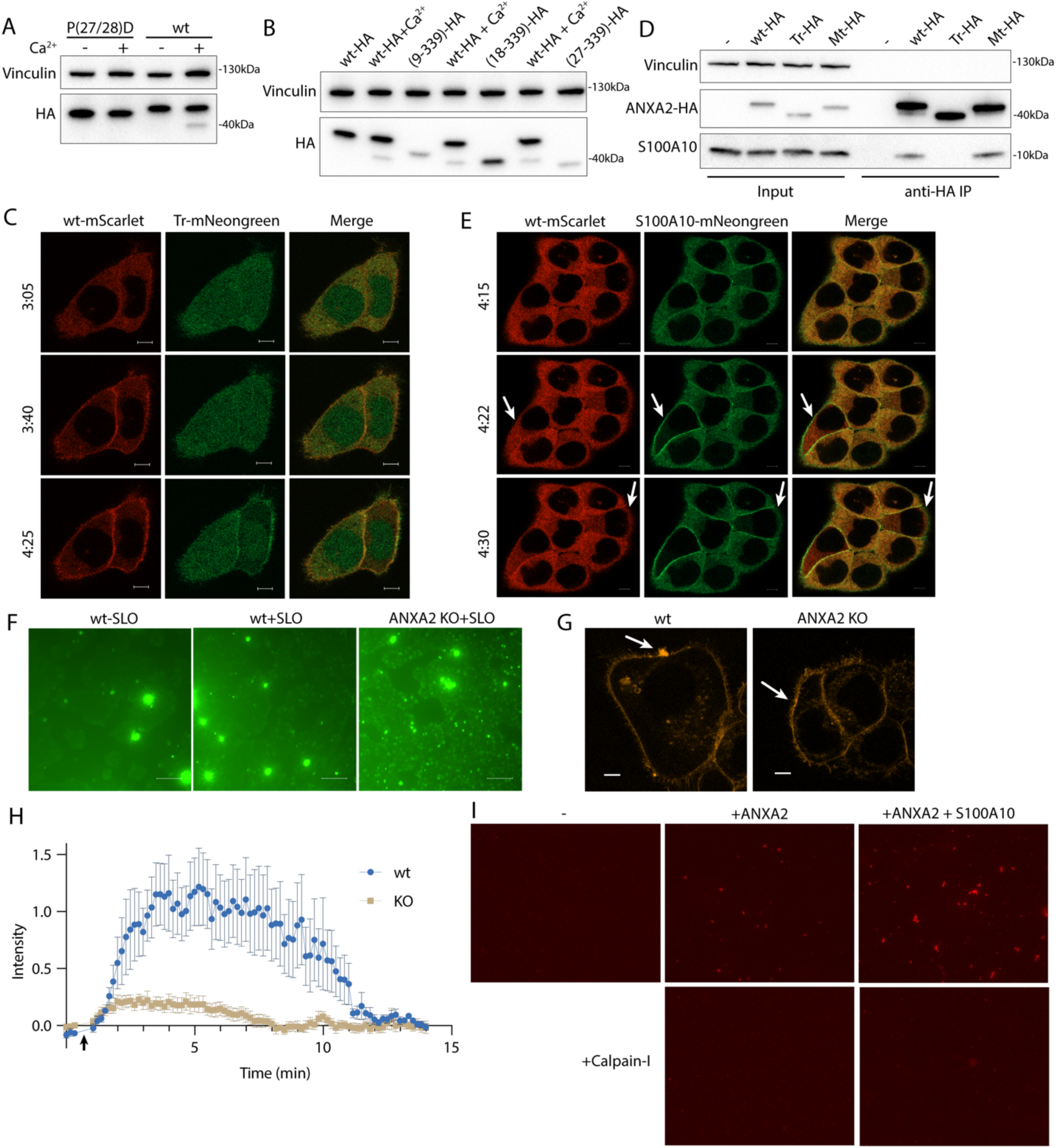
Calpain cleavage attenuates the membrane binding and scabbing activity of annexin A2. (A) Immunoblot analysis of lysates from cells expressing wildtype ANXA2-HA or ANXA2[P27D, P28D]-HA. Ca^2+^ (1 mM) was added to lysis buffer where indicated. (B) Immunoblot analysis of lysates from cells expressing wildtype ANXA2-HA or ANXA2-HA with the indicated N-terminal truncations. Ca^2+^ (1 mM) was added to lysis buffer where indicated. (C) Representative confocal micrographs of ANXA2-mScarlet (wt-mScarlet), ANXA2(18-339)-mNeonGreen (Tr-mNeonGreen)-expressing cells. Image times are relative to the addition of 400 ng/ml SLO. Scale Bars: 5 μm. (D) Immunoblot analysis of lysates and anti-HA immunoprecipitation elutions from cells expressing wildtype ANXA2-HA (wt-HA), ANXA2(18-339)-HA (Tr-HA), ANXA2[P27D, P28D]-HA (Mt-HA), or no HA construct. (E) Representative confocal micrographs of ANXA2-mScarlet (wt-mScarlet) or S100A10-mNeonGreen-expressing cells. Image times are relative to the addition of 400 ng/ml SLO. Arrows indicate cells with annexin A2 and S100A10 translocating to the membrane. Scale Bars: 5 μm. (F) Representative widefield micrographs of wildtype or annexin A2 knockout cells, with 2.5 μM Sytox Green added for 6 min. SLO (200 ng/ml) was added with Sytox Green in the indicated panels. Scale Bars: 100 μm. (G) Representative confocal micrographs of FM1-43 stained (2.5 μM) wildtype or annexin A2 knockout cells, 100 sec after laser ablation. Scale Bars: 5 μm. (H) Quantification over time of the repair scab intensity from FM1-43 stained (2.5 μM) wildtype or annexin A2 knockout cells. 6 cells were ablated for each condition. (I) Representative widefield micrographs of Texas Red-labeled 200 nm liposomes mixed with the indicated combinations of 300 nM annexin A2, annexin A2-S100A10, or Calpain-1.

Although cleaved annexin A2 retained some Ca^2+^-dependent membrane association activity *in vitro* (Fig. S5a), its membrane binding was attenuated (Figure 5c). High doses of SLO cause large scale translocation of annexins to the plasma membrane (34). After HCT116 cells were treated with a high, near-lytic (400 ng/ml) dose of SLO, tr-annexin A2-mNeonGreen was recruited more slowly to the plasma membrane compared to wt-annexin A2-mScarlet. In addition, a significant pool of tr-annexin A2 remained cytosolic unlike wt-annexin A2 which was almost completely recruited to the membrane. Mt-annexin A2 was recruited as quickly as wt-annexin A2 to the plasma membrane (Fig. S5b).

S100 proteins bind to and modify annexin function. The first 10-12 amino acid segment of annexin A2 binds to S100A10 (K_d_ = 13nM) (35). Thus, we predicted that calpain-cleaved annexin A2 would be incapable of interacting with S100A10. Immunoprecipitation of wt- and mt-annexin A2-HA captured S100A10. As expected, however, tr-annexin A2-HA no longer coprecipitated S100A10 (Figure 5d). Because previous reports suggest that annexin A2 may interact with other S100 proteins (36), we tested the specificity of the annexin A2-S100A10 interaction. Using annexin A2-Halo or calpain-treated annexin A2-Halo as bait, we captured annexin A2 interacting proteins (Fig. S5c). Two proteins of apparent molecular masses of 10 and 40 kDa appeared in the annexin A2-halo elution, but not in the calpain-treated annexin A2-halo elution. Gel excision-mass spectrometry identified these proteins as annexin A2 and S100A10, with no peptides mapping to any other S100 protein.

Annexin A2 and S100A10 formed an oligomer (Fig. S5f). We wished to test the possibility that this oligomer promoted the tight recruitment and membrane bridging activity of annexin A2. A reduced binding of S100A10 might account for the lowered membrane recruitment of tr-annexin A2. The membrane recruitment of S100A10 was enhanced in comparison to wt-annexin A2 after HCT116 cells were treated with a high dose of SLO (Figure 5e). We noted that S100A10 required annexin to bind membranes (Fig. S5d) and was destabilized and degraded *in vivo* after annexin A2 depletion (37). This suggested that all membrane recruited S100A10 is bound to a pool of annexin A2 and that calpain dissociates the S100A10-annexin A2 dimer of dimers.

We confirmed that annexin A2 is required for membrane repair in our HCT116 cell model. Using a CRISPR Cas9 intron trapping approach (38), we generated an annexin A2 knockout (Figure 6b). Scabbing/plugging was monitored over time in the presence of a high concentration of Sytox Green (2.5 μM) and SLO. After 6 min of SLO treatment, significantly more Sytox Green entered annexin A2 knockout cells compared to wildtype HCT116 cells (Figure 5f). This difference was dependent on addition of calcium to the medium (Fig. S5e). In a 37°C incubation, SLO binding and pore formation took closer to 3-4 min (Fig. S5e), suggesting that the increased Sytox Green influx in annexin A2 KO cells occurred within 2 min of pore formation. We next tested the effect of annexin A2 KO on repair scab formation (Figure 5g, 5h). Wild-type cells ablated with a laser in the presence of FM1-43 revealed a brightly stained repair scab distended from the ablation site by 1 minute post injury. This repair scab was nearly absent in annexin A2 KO cells. We conclude that annexin A2 knockout caused a defect in scabbing during the early stages of membrane repair.

**6.**
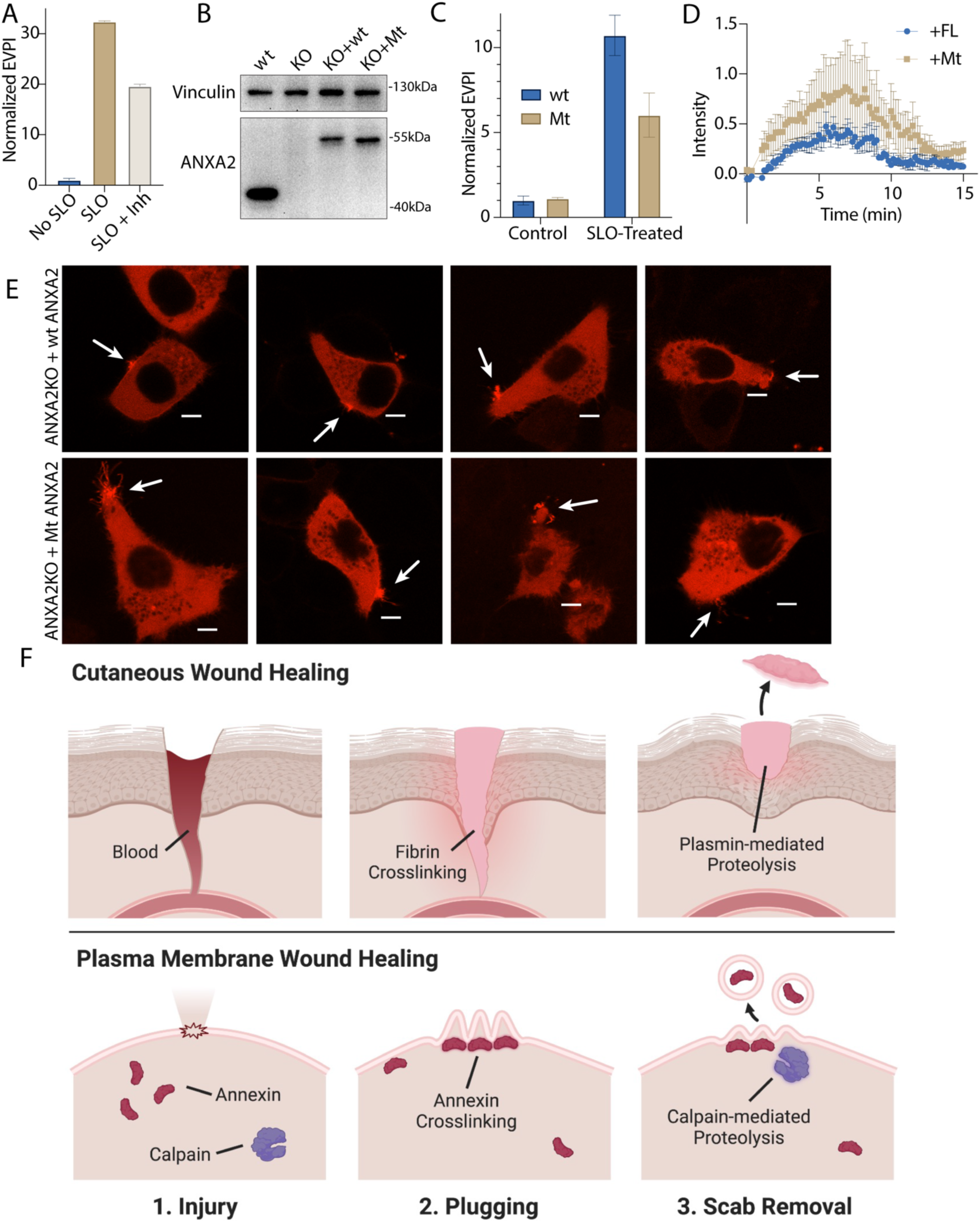
Inhibition of annexin A2 cleavage decreases annexin A2^+^ microvesicle secretion during membrane repair. (A) EV production index from ANXA2-Nluc cells treated with the indicated combinations of 200ng/ml SLO and 20 μM calpain Inhibitor, ALLN. (B) Immunoblot analysis of lysates from wildtype cells (wt), annexin A2 knockout cells (KO), and annexin A2 knockout cells expressing wildtype annexin A2-Nluc (KO + wt) or ANXA2[P27D, P28D]-Nluc (KO + Mt). (C) EV production index from wildtype ANXA2-Nluc (wt) or ANXA2[P27D, P28D]-Nluc (Mt) cells treated with or without SLO. (D) Quantification over time of the repair scab intensity from FM1-43 stained (2.5 μM) annexin A2 knockout cells rescued with ANXA2-mScarlet (wt) or ANXA2[P27D, P28D]-mScarlet (Mt). Cells (6) were ablated for each condition. (E) Confocal micrographs of 8 laser ablated annexin A2 knockout cells rescued with either wild-type ANXA2-mScarlet (wt) or ANXA2[P27D, P28D]-mScarlet (Mt). Images are 2 min, 30 sec after ablation. Scale Bars: 5 μm. (F) Schematic depicting the current model of plasma membrane repair and annexin^+^ MV secretion.

Annexin A2-S100A10 dimer of dimers aggregate liposomes *in vitro* (39) in a manner that may resemble lesion site scabbing *in vivo*. We purified both annexin A2 and annexin A2-S100A10 dimer of dimers (Fig. S5f) and mixed them with liposomes. Addition of annexin A2 or, to a larger extent, addition of annexin A2-S100A10 caused liposomes to aggregate (Figure 5i). Addition of calpain-1 dissolved annexin A2 and annexin A2-S100A10 liposome aggregates. Thus, calpain cleavage reduces annexin A2 membrane recruitment, terminates annexin A2-S100A10 association, and dissolves annexin A2-mediated membrane scabs.

The catalytic activity of the calpains is upregulated by PI_(4,5)_P_2_ allowing the proteins to partially associate with membranes in vivo (40). We wondered whether calpains, like annexins, could be recruited to membranes in the presence of calcium (Fig. S5G). We centrifuged membranes from the post nuclear supernatant (PNS) of a cell lysate, either in the presence or absence of 1 mM Ca^2+^. Membranes sedimented in the presence of Ca^2+^ were washed with 5 mM EGTA. High levels of annexin A1 and A2 and activated, autolyzed calpain-1 were eluted and recovered in the wash supernatant sample. In contrast, annexins and calpain-1 did not sediment along with membranes when 1 mM Ca^2+^ was not added to the PNS. Thus, calpains are also recruited to membranes in the presence of calcium Ca^2+^, either by directly binding membrane lipids or by interacting with membrane-associated proteins.

### Inhibiting annexin cleavage causes defects in repair scab secretion during membrane repair

We next tested the role of calpain-mediated cleavage in the shedding of annexin A2 scabs (Figure 6a). Calpain inhibition significantly lowered annexin A2^+^ MV secretion. To test the specificity of the effect to annexin A2, we transfected annexin A2 KO cells with wt-annexin A2-Nluc or mt-annexin A2-Nluc (Figure 6b). SLO-treated cells secreted lower levels of vesicular mt-annexin A2 compared to wt-annexin A2 (Figure 6c). Next, we transfected annexin A2 KO cells with wt-annexin A2-mScarlet or mt-annexin A2-mScarlet and imaged rescue after laser ablation (Figure 6d, 6e). Transfection of annexin A2-mScarlet partially rescued repair scab formation. Quantification of the FM1-43-stained repair scab revealed that cells rescued with mt-annexin A2-mScarlet had a larger repair scab compared with cells rescued with wt-annexin A2-mScarlet, with some overlap in the error bars during the time course (Figure 6d). Visualization of annexin A2-mScarlet after ablation revealed that the mt-annexin A2^+^ repair scab often had long (5 μm) filipodia-like structures extending from the repair site. In contrast, the mt-annexin A2^+^ repair scab was smaller and more compact. We conclude annexin cleavage is an integral part of the membrane repair and shedding process.

## Discussion

### Annexin-dependent membrane repair is a two-step process

We find that calpains target annexins during plasma membrane repair, driving secretion of damaged membranes as microvesicles and suggest a model of plasma membrane repair that is analogous to cutaneous wound repair (Figure 6f). After epidermal damage, blood flows into the wound and clots. Clotting is driven fibrin cross linking and scabbing over the wound. In a second, slower process, proteases including plasmin and collagenase cut the crosslinks, and the scab eventually falls off the repaired wound (41). During plasma membrane repair, annexins, including annexin A2, are rapidly recruited to the plasma membrane within 45 sec of damage induced by laser ablation (Figure 2b) or SLO (Figure 5c). An annexin^+^ membrane scab-like structure forms at the lesion site that may be visualized with the dye, FM1-43, and slows FM1-43 staining of intracellular organelles (Figure 2a). Like fibrin, annexin A2 and other annexins may mediate this scabbing by crosslinking membranes at the wound site. *In vitro*, annexins crosslink liposomes (Figure 5I). Annexin A2 knockout cells lose latency to a membrane impermeable dye within 2 min of wounding (Figure 5f), a period that coincides with annexin A2 recruitment to the membrane lesion (Figure 2b). Finally, the FM1-43-stained repair scab is virtually absent in laser-damage annexin A2 knockout cells (Figure 5g, 5h). In a second stage, as for plasmin, calpain-1/2 cleave annexins (Figure 3a, 3c, 4a-e). Calpain cleavage of annexin A2 peaks by 1-2 min and is spatially restricted within repair scabs (Figure 4h). Calpain cleavage of annexin A2 dissociates aggregated liposomes *in vitro* (Figure 5f) and prevents S100A10 binding (Figure 5d). Calpain cleavage and dissociation promotes secretion of annexin A2 in microvesicles (Figure 6a-c). Secretion of both perforated and closed vesicles (Fig. S3b-c) peaks by 5 min and is completed by 10 min (Figure 2d). Rescue of annexin A2 knockout with uncleavable annexin A2 causes a larger repair scab to form (Figure 6d, 6e). This descabbing process may increase the flexibility of the repair scab to facilitate budding and shedding, leading to a smaller repair scab at steady state. Annexins form blebs (42) and lattice-like sheets *in vitro* (43). Cleavage may also destabilize these structures, potentially preventing annexin overactivation.

Depending on the type of plasma membrane wound and the extent of damage, different repair machinery may be employed. While some reports suggest that calpains and annexins are important for pore toxin repair (12), other reports propose that this machinery is reserved for repair of larger lesions (13, 23). Although our data supports that annexins and calpains are important for repairing smaller SLO pores, this machinery may be even more important for repair of large lesions. Additionally, different cell lines may rely on different machinery.

### Substrate trapping reveals new targets for calpain-1/2

Using unbiased proteomics, we identified targets for calpain-1 and -2. Because calpain substrate recognition appears not to rely on primary sequence alone, computationally predicting calpain targets is challenging. Our dataset predicts hundreds of new calpain targets which may aid the development of new prediction algorithms. Although calpains were known to cleave membrane-cytoskeleton anchors during repair (44), we have extended the targets to include annexins A1, A2 and A6 (Figure 4i-j, Figure 4b, S4d). All three annexins aggregate liposomes in a manner that is reversed by calpain cleavage of annexin A2 (Figure 5f), and likely also for annexin A1 and A6. Annexin A2 is thought to form a “membrane repair complex” with S100A10, AHNAK, and with dysferlin in muscle cells (32). This complex may mediate calcium-dependent patching of membrane lesions. AHNAK and S100A10 were identified in our proteomic analysis of annexin A2 binding partners. In muscle cells, dysferlin interacts with annexin A2 and is also directly targeted by calpain 1/2 (45). We suggest that by targeting annexins, AHNAK and dysferlin, calpains may control membrane scabbing proteins during calcium-dependent membrane repair. Low level expression of a C-terminal truncation mutant of annexin A6, similar to the calpain-cleaved form of annexin A6, caused dominant-negative inhibition of membrane repair (7, 18). This observation supports our view that annexin cleavage destabilizes the repair cap and likely occurs after scrabbing. If calpains cleave annexin A6 prior to annexin recruitment, scrabbing/repair cap formation is inhibited.

Calpain cleavage may modulate aspects of annexin function other than cell surface scabbing and secretion of MVs. The N-terminal domain of annexins interact with other proteins directly or indirectly through S100 proteins. Annexin A2, for example, may interact with the cytoskeleton (12, 36). Calpain cleavage would terminate such an interaction. Both calpain activities, descabbing of annexins and detaching of the membrane from the cytoskeleton may be important for membrane repair and shedding.

### Ca^2+^-dependent production of MVs secrete annexins and annexin cleavage products

Lower levels of annexin-containing microvesicles are secreted into the culture medium by unstimulated cells (Figure 1a-c). It is tempting to speculate that some low level of membrane damage, repair and MV secretion occurs continuously. Consistent with this model, MVs are enriched with an array of Ca^2+^-responsive proteins (Figure 1e), many of which are important for membrane repair. Also, the annexins in constitutively secreted MVs are cleaved by calpains, which requires micromolar calcium influx for activation. Such “damage” may be a consequence of the passaging of cells or the use of bovine serum which contains complement proteins capable of perforating human cells (46). Other processes such as cell death or the activation of a very high flux channel (e.g. P2X7) may elevate Ca^2+^ to levels sufficient for secretion of these vesicles (47). We suggest that these are normal processes that occur in animals and may explain the representation of MVs in all bodily fluids.

N-terminal peptides from annexins, particularly from annexin A1, mediate anti-inflammatory signaling. Annexin A1-derived peptides bind to Formyl Peptide Receptors (FPRs) on the surface of immune cells, dampening proinflammatory signals and promoting proresolving processes such as efferocytosis (48). We propose that calpains may also generate these peptides. Once cleaved, N-terminal peptides may diffuse directly through the membrane or through vesicular disruptions. Annexin A1 N-terminal peptide intercalates into phosopholid membranes, suggesting that it may penetrate membranes (49). These peptides could then stimulate immunologically silent disposal of damaged membrane products.

## Methods

### Cell lines, media, general chemicals, and DNA

HEK293T, C2C12 HCT116, and all HCT116-derived cell lines were grown at 37°C in 5% CO_2_ in Dulbecco’s Modified Eagle’s Medium (DMEM) supplemented with 10% fetal bovine serum (FBS) (Thermo Fisher Scientific, Waltham, MA, USA). Cells were routinely tested, and found negative, for mycoplasma contamination with the MycoAlert Mycoplasma Detection Kit (Lonza Biosciences). HCT116, C2C12, and HEK293T cells were authenticated by the UC Berkeley Cell Culture Facility using STR profiling. For Figures 1 and S1, HCT116 or HEK293T cells were incubated in EV-depleted medium produced by ultracentrifugation of DMEM supplemented with 25% FBS at 186,000 × g (40,000 RPM) for 24 h using a Type 45Ti rotor. Ultracentrifuged media was then diluted to 10% FBS with DMEM. For SILAC experiments, HEK293T cells were grown in High Glucose DMEM for SILAC (Athena Enzyme Systems), supplemented with 10% dialyzed FBS (Thermo Fisher Scientific, Waltham, MA, USA) for 2 wk prior to exosome isolation. DMEM was supplemented with GlutaMax, L-leucine (800 μM), L-methionine (200 μM), and either L-lysine (800 μM) and L-arginine (400 μM) or L-lysine ^13^C_6_, ^15^N_2_ (800 μM) and L-arginine ^13^C_6_, ^15^N_4_ (400 μM). EV-depleted SILAC medium was prepared as described for EV-depletion of normal medium. C2C12 cells were differentiated for 5 d in DMEM supplemented with 2% horse serum and 1 μM insulin. Antibodies for immunoblot were: rabbit anti-vinculin (Abcam ab129002), rabbit anti-annexin A6 (Abcam 201024), rabbit anti-annexin A1 (Abcam ab214486), rabbit anti-annexin A2 (Abcam ab185957), rabbit anti-calpain 1 (Proteintech 10538-1-AP), mouse anti-CD63 (BD Biosciences 556019), rat anti-CD63 (LSBio LS-C179520), mouse anti-alix (Santa Cruz sc-53540), mouse anti-flotillin-2 (BD Biosciences 610384), rabbit anti-CD9 (CST D801A), mouse anti-tubulin (Abcam ab7291), rabbit anti-streptolysin O (GeneTex GTX64171), mouse anti-NanoLuc (R&D Systems MAB10026), rabbit anti-HA (CST C29F4), rabbit anti-LC3 (Novus NB100-2220), rabbit anti-Lamp1 (CST D2D11).

### Lentivirus production and transduction

The pLJM1 backbone was used for stable expression of annexin A2 and S100A10 in HCT116 cells. Fusion protein domains were separated by (GGSG)_2_ linkers. For low expression of annexin A2-Nluc (Figure 2, S2, S3), wt/Tr/Mt-annexin A2-mScarlet/mNeonGreen (Figure 5, S5), wt/Tr/Mt-annexin A2-HA (Figure 5) and S100A10-mNeonGreen (Figure 5), the pLJM1 CMV promoter was replaced with the RPL30 promoter. pLJM1 plasmids expressing mNeonGreen-tagged proteins contained puromycin resistance cassettes. For pLJM1 plasmids expressing mScarlet and Nluc-tagged proteins, the puromycin resistance cassette was replaced with a blasticidin resistance cassette. The pLIX403-puro backbone was used for stable, doxycycline-inducible mCherry-VPS4a(E228Q) expression.

HEK293T cells at 40% confluence within a 6-well plate were transfected with 165 ng of pMD2.G, 1.35 µg of psPAX2, and 1.5 µg of a pLJM1 or pLIX403 plasmid using the TransIT-LT1 Transfection Reagent (Mirus Bio) as per the manufacturer’s protocol. At 48 h post-transfection, 1 ml of fresh DMEM supplemented with 10% FBS was added to each well. The lentivirus-containing medium was harvested 72 h post-transfection by filtration through a 0.45 μm polyethersulfone (PES) filter (VWR Sciences). The filtered lentivirus was distributed in aliquots, snap-frozen in liquid nitrogen, and stored at –80°C. For lentiviral transductions, we infected HCT116 cells with filtered lentivirus in the presence of 8 μg/ml polybrene for 24 h, and the medium was replaced. HCT116 cells were selected using 1 μg/ml puromycin or 4 μg/ml blasticidin S for 4 days and 6 d, respectively. Gene expression was assessed by immunoblot analysis.

### Immunoblotting

Cells were washed once with PBS and lysed in TBS containing 1% TX-100, 1 mM EGTA, and a protease inhibitor cocktail (1 mM 4-aminobenzamidine dihydrochloride, 1 µg/ml antipain dihydrochloride, 1 µg/ml aprotinin, 1 µg/ml leupeptin, 1 µg/ml chymostatin, 1 mM phenylmethylsulfonyl fluoride, 50 µM N-tosyl-L-phenylalanine chloromethyl ketone, and 1 µg/ml pepstatin), and incubated on ice for 10 min. The whole cell lysate was centrifuged at 1,000 × g for 10 min at 4°C and the post-nuclear supernatant was diluted with 6× Laemmli buffer (without DTT) to a 1× final concentration. Samples were heated at 95°C for 5 min and proteins resolved on 4–20% acrylamide Tris-glycine gradient gels (Life Technologies). Proteins were then transferred to polyvinylidene difluoride membranes (EMD Millipore, Darmstadt, Germany), blocked with 5% bovine serum albumin (BSA) in TBS, washed 3× with TBS-T and incubated overnight with primary antibodies in 5% BSA in TBS-T. The membranes were then washed again 3× with TBS-T, incubated for 1 h at room temperature with 1:10,000 dilutions of anti-rabbit or anti-mouse secondary antibodies (GE Healthcare Life Sciences, Pittsburgh, PA, USA), washed 3× with TBS-T, washed once with TBS and then detected with ECL-2 or PicoPLUS reagents (Thermo Fisher Scientific) for proteins from cell lysates or EV isolations, respectively.

### Synchronized SLO Treatment

Lyophilized SLO (Sigma) was pre-activated in PBS containing 10 mM DTT at 37°C for 1 h, distributed in aliquots into low-retention microcentrifuge tubes, snap-frozen in liquid nitrogen, and stored at –80°C until use. The protein concentration of the SLO batch was determined by a Bradford assay. HCT116 cells plated two days before (70-80% confluent) were washed with cold PBS and incubated in prechilled DMEM (w/o Calcium), 1 mM EGTA, and the indicated SLO concentration for 15 min at 4°C. Cells were washed once with cold PBS and replaced with prewarmed 37°C DMEM (w/o Calcium) + 1.8 mM CaCl_2_ and incubated for 20 min at 37°C.

### Crude, High-Speed Extracellular Vesicle Pellet Centrifugation

Conditioned medium from 8 x 15 cm plates (240 ml) was harvested from untreated (Figure 1) or 200 ng/ml SLO-treated (Fig. S2e, 3a, 3b, 4e) HCT116 cells. All subsequent manipulations were completed at 4°C. Cells and large debris were removed by low-speed sedimentation at 1,000 × g for 15 min in a Sorvall R6 +centrifuge (Thermo Fisher Scientific) followed by medium-speed sedimentation at 10,000 × g for 15 min using a fixed angle FIBERlite F14−6×500 y rotor (Thermo Fisher Scientific). The supernatant fraction was then centrifuged at 29,500 RPM (∼100k x g) for 1.25 h in a SW32 rotor. The high-speed pellet fractions were resuspended in PBS unless otherwise stated and pooled (600 μl total).

### EV fractionation by buoyant iodixanol density gradient equilibration

Fresh aliquots of 5.4, 10.8, 16.2, and 21.6% (v/v) iodixanol solutions were prepared by mixing appropriate volumes of PBS and Solution D (PBS and 54% (w/v) iodixanol). A 27% (v/v) iodixanol solution was prepared by mixing the resuspended high-speed pellet with Solution D. Iodixanol gradients were prepared by sequential 1 ml overlays of each iodixanol solution in a 5 ml SW55 tube, starting with the 27% iodixanol solution and finishing with the 5.4% iodixanol solution. Gradients were centrifuged in a SW55 rotor at 36,500 RPM for 16 h with acceleration set at a minimum level and no brake. Fractions (350 µl) were collected from top to bottom. Density measurements were taken using a refractometer. Each fraction was diluted in 6× Laemmli buffer (without DTT) for immunoblot analysis.

### EV fractionation by buoyant sucrose step gradient equilibration

Fresh aliquots of 10, 40, and 60% (v/v) sucrose solutions were prepared in 20 mM Tris (pH 7.4). A 55% (v/v) sucrose solution was prepared by mixing the resuspended high-speed pellet with the 60% sucrose solution. Sucrose gradients were prepared by sequentially layering 1 ml of the 40% and 10% sucrose solution over 3 ml of the 55% sucrose solution in a 5 ml SW55 tube. Gradients were centrifuged in a SW55 rotor at 36,500 RPM for 16 h with minimum acceleration and no brake. Fractions (400 µl) were collected from top to bottom. Density measurements were taken using a refractometer. Each fraction was diluted in 6× Laemmli buffer (without DTT) for immunoblot analysis.

For SILAC experiments, after flotation, the 3rd and 4th fraction from the top of gradient were diluted with 10 ml PBS and centrifuged to sediment EVs. Pellet fractions were resuspended in TBS + 2% SDS, and protein concentration was determined using a microBCA kit (Thermo). Fraction 3 protein (1 µg) from heavy cells was mixed with Fraction 4 protein (1 µg) from light cells and vice versa. Glycerol (10%) was added to both mixes and samples were loaded onto a SDS page gel. Proteins were electrophoresed into the stacking gel, and the gel was stained using Sypro Ruby using the overnight protocol (Thermo). Protein samples were excised from the gel, digested in-gel with trypsin, and analyzed by quantitative mass spectrometry.

### Exosome Pulldowns

For CD63^+^ EV immunoprecipitation, Protein G Magnetic Dynabeads (50 µl) (Thermo Fisher Scientific) were washed twice with 200 µl PBS using a magnetic rack to capture beads between washes. The resuspended high-speed pellet fraction and 1 μg anti-CD63 (clone H5C6, BD Biosciences), was added to the beads and incubated with rotation overnight at 4°C. The beads were washed 3x with 600 µl cold PBS, incubating for 1 min between washes. EVs were eluted with 30 µl PBS + 0.1% TritonX-100 for 5 min. Fractions were diluted in 6× Laemmli buffer (without DTT) for immunoblot analysis.

For pulldown of EVs with exposed phospholipids using annexin A5, the high-speed extracellular vesicle pellet fraction from 2 x 15 cm plates of SLO-treated cells was resuspended in 200 ul of annexin binding buffer (10 mM HEPES pH 7.4, 140 mM NaCl, 2mM calcium chloride). Biotin-X ANXA5 (5 µl)(Thermo Fisher Scientific) was added, mixed, and incubated for 15 min at room temperature. Annexin binding buffer (350 µl) and 50 µl of Steptavidin Dynabeads (Thermo Fisher Scientific), prewashed with 50 µl annexin binding buffer, was added to the binding reaction and incubated for 10 minutes. The beads were washed twice with 400 µl annexin binding buffer and eluted with 200 µl of 10 mM HEPES pH 7.4, 140 mM NaCl, 2mM EGTA, 0.1% TX-100. Fractions were diluted in 6× Laemmli buffer (without DTT) for immunoblot analysis.

### ANXA2-Nluc and CD63-Nluc secretion assay

Assays were performed as described previously (20), with slight modifications. HCT116 CD63-Nluc or ANXA2-Nluc cells were grown to ∼80% confluence in 24-well plates. All subsequent manipulations were performed at 4°C. Conditioned medium (200 µl) was taken from the appropriate wells, added to a microcentrifuge tube, and centrifuged at 1000 × g for 15 min in an Eppendorf 5430 R centrifuge (Eppendorf, Hamburg, Germany) to remove intact cells. Supernatant fractions (150 µl) from the low-speed sedimentation were passed through a 0.45-micron filter (96-well format) by centrifuging at 1000 × g for 5 min. Filtered fractions (50 µl) were used to measure luminescence. During these centrifugation steps, the cells were placed on ice, washed once with cold PBS, and lysed in 200 µl of PBS containing 1% TX-100 and protease inhibitor cocktail.

To measure vesicular Nluc secretion, we prepared a master mix containing the membrane-permeable Nluc substrate and a membrane-impermeable Nluc inhibitor using a 1:1000 dilution of Extracellular NanoLuc Inhibitor and a 1:333 dilution of NanoBRET Nano-Glo Substrate into PBS (Promega, Madison, WI, USA). Aliquots of the Nluc substrate/inhibitor master mix (100 µl) were added to 50 µl of the supernatant fraction from the medium-speed centrifugation and vortexed briefly, and luminescence was measured using a Promega GlowMax 20/20 Luminometer (Promega, Madison, WI, USA). For the intracellular normalization measurement, the luminescence of 50 µl of cell lysate was measured using the Nano-Glo Luciferase Assay kit (Promega, Madison, WI, USA) as per the manufacturer’s protocol. The exosome production index (EPI) for each sample is calculated as follows: EPI = medium/cell lysate.

### Isolation of cytosol from cultured human cells

Isolations were performed as described previously (20), with slight modifications. HCT116 WT cells were grown to ∼90% confluence in 150 mm dishes. All subsequent manipulations were performed at 4°C. Each 150 mm dish was washed once with 10 ml of cold PBS and then harvested by scraping into 5 ml of cold PBS + 1 mM EGTA. The 5 ml cell suspensions from either 4 (Figure 4a, 4b) or 20 (Figure 4i) 150 mm dishes were combined. Cells were then collected by centrifugation at 200 × g for 5 min, and the supernatant fraction was discarded. The cell pellet was resuspended in 2 vol of lysis buffer (TBS, 1 mM EGTA, 0.2 mM PMSF, and for Figure 4i, 10 µM E64) and placed on ice. Cells were mechanically lysed by 15 strokes through a 22-gauge needle. Cell lysates were centrifuged at 1000 × g for 15 min to sediment intact cells and nuclei, and the post-nuclear supernatant was then centrifuged at 49,000 RPM for 15 min in an TLA-55 ultracentrifuge (Beckman Coulter). The supernatant (cytosol fraction) was collected conservatively without disturbing the pellet.

### Protein purification of ANXA2, ANXA2-S100A10 Complex, ANXA2-HaloTag

Pet28a vectors expressing ANXA2, S100A10, or ANXA2-HaloTag were transformed into Rosetta (DE3) BL21 *E. coli*, and grown in 250 ml LB cultures at 37°C. At O.D. = 0.5, protein expression was induced with 50 µM IPTG. Cultures were grown overnight at 18°C and centrifuged at 3k x g for 10 min. Pellet fractions were kept at −80°C until time for purification. Pellet samples were thawed and resuspended in 7.5 ml *E. coli* Lysis Buffer (1x TBS, 1x Protease Inhibitor Cocktail, 10 μl Benzonase [NEB], 1 mg/ml lysozyme, 5 mM DTT, 1 mM EGTA). Bacteria were lysed by sonication for 5 sec on (20% power), 20 sec off, 5 times. *E. coli* lysate was centrifuged for 5 min at 1,000 x g at 4°C. The supernatant fraction was removed and transferred to a high-speed, 1.5 ml centrifuge tube (Beckman Coulter). For purification of ANXA2-S100A10 complex only, ANXA2 lysate and S100A10 lysate were mixed 1:1. 2 mM CaCl_2_ (final) was added and incubated for 5 min at room temperature. Lysates were centrifuged at >100k x g (49k RPM) in a TLA 55 for 10 min and the supernatant was discarded. The membrane pellet was resuspended with a 5 ml annexin wash buffer (1x TBS, 5 mM CaCl_2_, 5 mM ATP, 5 mM DTT), using a 25-gauge needle for thorough resuspension. The resuspended pellet was centrifuged again at 100k x g (49k RPM) in a TLA 55 rotor for 10 min. The supernatant fraction was carefully removed, and the membrane pellet resuspended with 1 ml annexin elution buffer (1x TBS, 5 mM DTT, 10 mM ATP). DTT was replaced with TCEP for downstream labeling of the eluted annexin A2 with quencher. The resuspended pellet fraction was centrifuged again at 100k x g (49k RPM) in a TLA 55 for 10 min. The supernatant fraction was carefully removed, and aliquots were flash frozen in liquid nitrogen for storage at −80°C. For labeling, annexin A2-Halo (14 µM) elution was mixed with 16 µM JF646 Halo Ligand (Promega). For self-quenched annexin A2-Halo, 286 µM Tide Quencher 5WS-Maleamide (AAT Bioquest) was also added to this reaction and incubated overnight at 4°C. The maleamide reaction was quenched with 5 mM DTT and unreacted Halo Ligand and Quencher were removed using two Bio-Spin 6 (Tris) columns (Bio-rad).

### Protein purification of dCAPN1 and dCAPN2

Human dCAPN1[C115S]-3xFlag-6xHis and dCAPN2[C105S]-3xFlag-6xHis were both cloned into PetDuet-1 coexpressing CAPNS1(86-269). *E. coli* cultures were grown at 37°C in 200 ml cultures to an O.D.600 of 0.5, and expression induced with 50 uM IPTG. Cultures were grown overnight at 18°C. *E. coli* were centrifuged at 3k x g for 10 min and resuspended in 5ml TBS + 0.2 mM PMSF. Cells were sonicated 5 times (5 sec on 15 sec off, 20% power), and lysates sedimented at 10k x g for 15 minutes. Supernatant fractions were applied to HisPure beads (500 μl of slurry) pre-washed with lysis buffer. Slurries were rotated at 4°C for 1 h and then washed 3 x with 5 ml 1.5x TBS + 20mM imidazole and centrifuged at 600 x g after washes to collect beads. Protein was eluted with 1 ml 1x TBS + 300mM imidazole. Aliquots (50 μl) were snap frozen in liquid nitrogen.

### Capture of CAPN1 and CAPN2 Substrates and Interactors

CaCl_2_ (2mM final) was added to E64-containing cytosol isolated as described above and incubated for approximately 1 h. In parallel, 125 μl anti-Flag agarose beads (Chromotek) were sedimented at 1k x g for 2 min and resuspended in 300 μl wash buffer (TBS + 1mM CaCl_2_ + 0.01% Tween-20). The resuspended beads were split into 3 tubes. Flag (3x) peptide, dCAPN1-3xFlag, or dCAPN2-3xFlag bait were added to the beads (6 μM final for each bait) in a 300 μl reaction. Bait proteins were bound to beads for 1 h at 4°C. Beads were wash once with 500 μl wash buffer and once with 500 μl lysis buffer (TBS + 10μM E64 + 0.2mM PMSF + 1 mM EGTA), with centrifugation at 1k x g for 2 min between washes to sediment beads. CaCl_2_-containing lysate (1.5 ml) was added to each of the 3 conditions and incubated for 2 h at 4°C. Beads were washed 3 with 1ml wash buffer, with centrifugation at 1k x g for 2 min between washes to sediment beads. Calpain substrates were eluted with 100 μl elution 1 buffer (TBS + 5mM EGTA), and stable calpain interacting proteins were eluted with 100 μl elution 2 (TBS + 1x Laemmli Buffer).

### Measurement of plasma membrane permeabilization by SLO over time

For measurement of plasma membrane permeabilization in bulk, we resuspended dissociated HCT116 cells in PBS + 5 mM EGTA. Cell slurries (50 ul, 4 x 10^5^ cells per reaction) were incubated on ice in a qPCR plate with the indicated concentration of SLO. Cells were sedimented at 300 x g and resuspended in 50 μl of ice-cold PBS + 5 mM EGTA + 2.5 μM SYTOX Green (Thermo Fisher). The plate was sealed with optical film and placed in a qPCR machine (Bio-Rad CFX96) pre-cooled to 4°C and then rapidly heated to 37°C with fluorescence measured every 20 sec using SYBR green settings. For measurement of plasma membrane permeabilization with microscopy, we grew HCT116 cells to 70% confluence in a 35 mm glass bottom dish (MatTek). Cells were placed in a 37°C, 5% CO_2_ chamber on an LSM900 microscope and then washed once with 1 ml Ca^2+^-free DMEM and incubated with Ca^2+^-free DMEM + 200 ng/ml SLO + 2.5 μM SYTOX Green. Where indicated, 1 mM Ca^2+^ was added to the incubation media. SYTOX Green uptake was measured by imaging with the 10x objective, taking images every 10 sec for 10 min.

### GUVs and Liposomes

For GUV assays, we mixed lipids in the following molar ratios: 38.5:20:20:3:18.5 DOPC:POPE:DOPS:PIP_(4,5)_P2:cholesterol (1 μmole total lipid) in 500 μl of 5% methanol, 95% chloroform. The lipid mix was smeared onto two indium tin oxide-coated glass plates, and for 30 min solvent was evaporated under vacuum on a heat block preheated to 55 °C. A circular rubber spacer coated with vacuum grease was placed on the glass slide into which buffer was introduced (1mM Tris pH 7.4, 250 mM sucrose), followed by covering with a second glass slide. Electrodes were clipped to each plate, and the chamber was placed in a 50°C oven. A function generator was used to apply an electric field (10 Hz, 1.4 V) for 90 min. For the last 30 min, the frequency was decreased by 0.5 Hz every 5 min. The GUV-containing solution was drained from the chamber and mixed with GUV buffer (5 mM Tris pH 7.4, 250 mM glucose). After settling to the bottom of the slide overnight at 4°C, GUVs were resuspended in GUV buffer with 2 mM CaCl_2_ and 5 mM TCEP and mixed with combinations of 1 μM ANXA2-Halo-JF646 and 1 μM S100A10-mScalet.

For liposome assays, we mixed lipids in a glass beaker in the following molar ratios: 38:20:20:3:0.5:18.5 DOPC:POPE:DOPS:PIP_(4,5)_P2:TexasRed-PE:cholesterol (1 umol total lipid) in 500 ul of 5% methanol, 95% chloroform. The lipid mix was placed on a heat block preheated to 55 °C and solvent was evaporated under vacuum for 30 min. Lipids were resuspended in degassed TBS and incubated at 55 °C with intermittent vortexing for 15 min. The lipid solution was pumped 15 times through an extruder with a 200-micron filter. Liposomes were diluted 1:5 into TBS with 2 mM CaCl_2_ and 5 mM TCEP, with indicated combinations of 300 nM untagged ANXA2, ANXA2-S100A10 complex, and porcine CAPN-I.

### Microscopy and Laser Ablation

The images in Figure 5g, S2c, S5d, were acquired on an Echo Revolve Microscope using the 10× air objective. The images in Figure 2a, 2b, S2b, 4h, 5c, 5e, s5b, and 6d were acquired using an LSM900 confocal microscope system (ZEISS) using confocal mode, a 63× Plan-Apochromat, NA 1.40 objective, and a heated, CO_2_ controlled chamber. Cells were bathed in DMEM with 2.5 µM FM1-43 (Biotium) added where indicated. For laser ablations using the LSM900, a 1 µM x 1 µM square was positioned over the edge of a cell not adjacent to another cell. The 1 µM x 1 µM square was ablated for 100-200 iterations using the UV laser at 100% power. Repair caps were quantified in Zen 3.1 (Zeiss) by measuring intensities within a box that encapsulated the repair cap. This intensity was normalized to a boxed piece of membrane that was not ablated.

## Supplementary Material

Video 1 shows shedding from the FM1-43-stained scab. Image times are relative to the first image taken post ablation. Arrow indicates the site of ablation. Scale Bars: 5 μm.

Table S1 provides ratios and p-values for enrichment of EV proteins in the low vs high buoyant density fractions of a sucrose gradient. Note that proteins detected only in one of the density fractions were eliminated.

Table S2 lists peptides detected from gel-excised annexin A2 protein treated with (Tab 2) or without (Tab 1) porcine calpain-1.

Table S3 provides protein abundances, calculated by peptide count, normalized spectral abundance factor (NSAF) or exponentially modified protein abundance index (emPAI) for EGTA elutions from pulldown reactions using 3xFlag, 3x-Flag C115S calpain-1 (CAPN1), or 3x-Flag C105S calpain-2 (CAPN2) as bait.

## Acknowledgments

We dedicate this work to Bob Lesch, our lab manager for the past several decades who was tragically taken from us by an accident in 2021. We would also like to thank our current lab manager, Nam Che, and the staff at the UC Berkeley shared facilities, the Cell Culture Facility (Alison Killilea) and the DNA Sequencing Facility. The mCherry-VPS4a (dominant mutant) cDNA was a gift from Kevin Rose in the laboratory of James H Hurley (UC Berkeley). This work used the Vincent J. Proteomics/Mass Spectrometry Laboratory at UC Berkeley, supported in part by NIH S10 Instrumentation Grant S10RR025622. JMN is supported by a National Institutes of Health F31 grant. RS is an Investigator of the Howard Hughes Medical Institute, a Senior Fellow of the UC Berkeley Miller Institute of Science, and Chair of the SAB of Aligning Science Across Parkinson’s Disease (ASAP). This work was funded by the Howard Hughes Medical Institute. The funders had no role in study design, data collection and interpretation, or the decision to submit the work for publication.

---

**S1.**
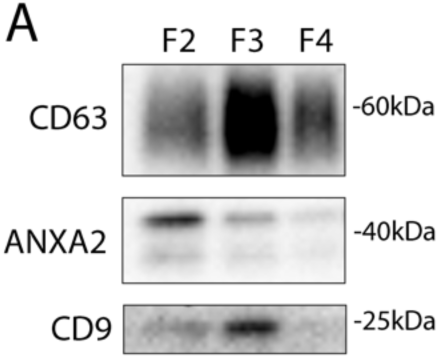
Annexin-containing extracellular vesicles are distinct from exosomes. Immunoblots show distribution of EV markers across a sucrose step gradient of the conditioned medium 100k × g pellet fraction. Samples taken from low density (F2-Fraction #2) to high density (F4-Fraction #4).

**S2.**
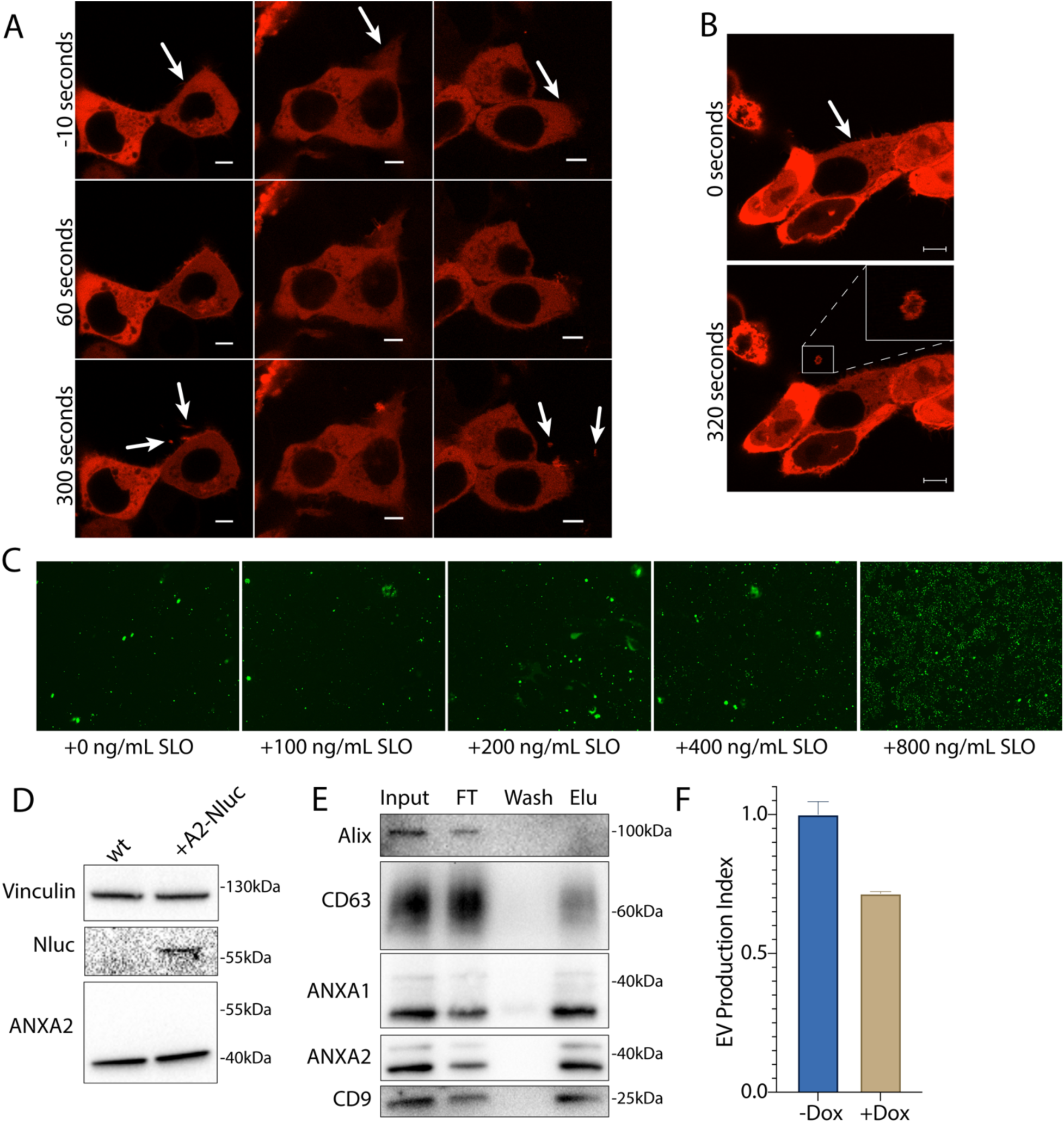
Annexin-containing extracellular vesicles are shed from the repair scab after plasma membrane damage. (A) Confocal micrographs of ANXA2-mScarlet recruitment in three laser ablation experiments are shown. Image times are relative to the first image taken post ablation. White arrows in the top panes indicate the sites of ablation. Arrows in the bottom panes indicate extracellular vesicles. Scale Bars: 5 μm. (B) Representative confocal micrographs of ANXA2-mScarlet shedding are shown. Image times are relative to the first image taken post ablation. White arrows in pane I indicate the site of ablation. Scale Bars: 5 μm. (C) Representative widefield micrographs of cells stained with 1 μM Sytox Green after a treatment period with the indicated SLO concentration and recovery period. (D) Immunoblots show expression of annexin A2-Nluc (A2-Nluc) using a low expression promoter. (E) Immunoblots show enrichment of EV markers after capture with immobilized annexin A5 from the conditioned medium 100k × g pellet fraction (FT-Flow Through, Elu-Elution). (F) EV production index from ANXA2-Nluc cells expressing mCherry-VPS4a (dominant mutant) under control of a doxycycline-inducible promoter. Cells were pretreated with 200ng/ml doxycycline (Dox) or DMSO for 6 h followed by treatment with 200 ng/ml SLO.

**S3.**
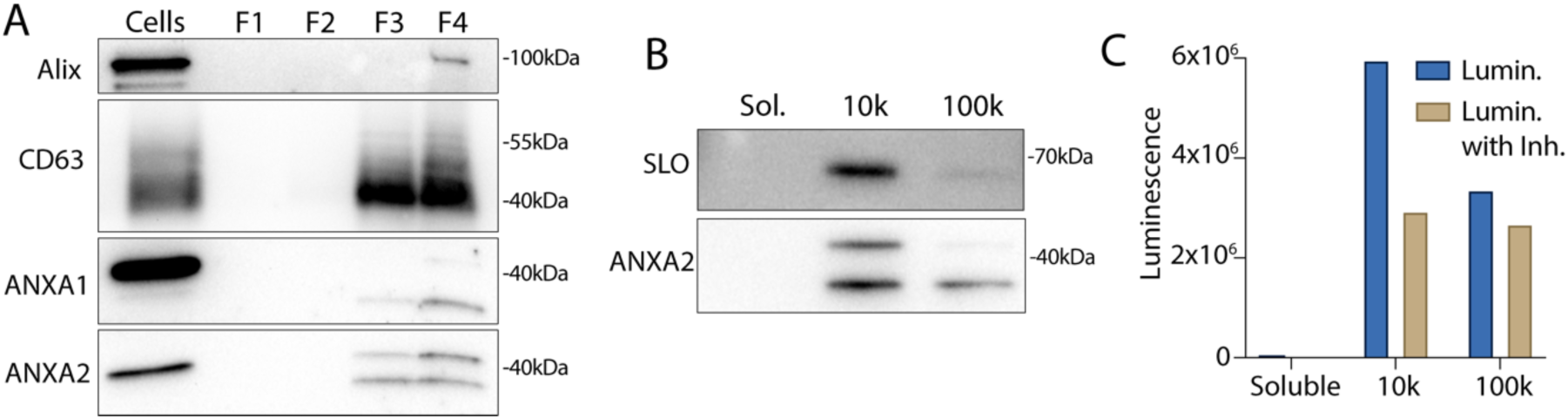
Annexins within extracellular vesicles are shifted in apparent molecular weight. (A) Immunoblots show enrichment of indicated EV markers relative to cell lysate after treatment of C2C12 myotubes with 200 ng/mL SLO. F1, F2, F3, and F4 refer to buoyant fractions of a sucrose step gradient of the conditioned medium 100k × g pellet fraction, moving from low to high density. (B) Immunoblot and (C) luminescence analysis of the 10k × g pellet fraction (10k), the 100k × g pellet fraction (100k), and the remaining soluble supernatant (Sol.) after serial centrifugation of conditioned media from ANXA2-Nluc cells treated with 200 ng/ml SLO. For each fraction, Nluc luminescence (lumin.) was measured with or without membrane impermeable Nluc inhibitor (Inh.).

**S4.**
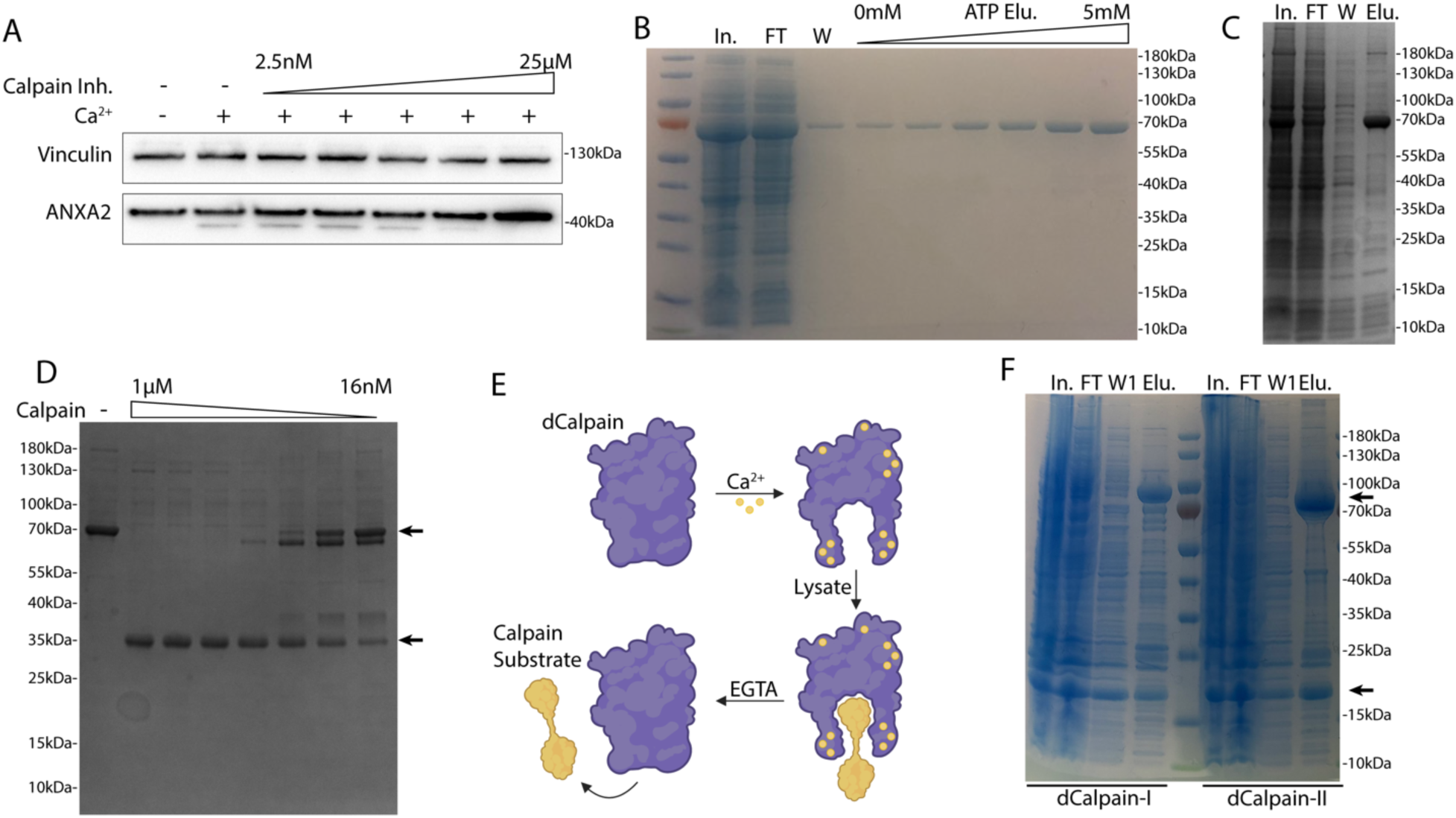
Calpain-1/2 cleave annexins which are then shed in microvesicles. (A) Immunoblot analysis of cytosol fractions after incubation with or without 1 mM Ca^2+^, with a range of concentrations of ALLN inhibitor (Calpain Inh.). (B) Coomassie-stained gel showing purification of annexin A2-Halo from *E. coli* (In- lysate input, FT- 100k x g pellet fraction flow through, W- CaCl_2_-containing 100k x g pellet wash, ATP Elu- elution from 100k x g pellet fraction with increasing concentrations of ATP). (C) Coomassie-stained gel showing purification of annexin A6-HA from *E. coli* (In- lysate input, FT- 100k x g pellet flow through, W- CaCl_2_-containing 100k x g pellet wash, Elu- elution off 100k x g pellet with 10mM ATP). (D) Coomassie-stained gel showing the mobility of recombinant annexin A6-HA, incubated with a range of concentrations of purified, porcine calpain-1. Arrows indicate uncleaved and cleaved product. (E) Schematic illustrating substrate binding and elution of substrates (yellow) to catalytic cysteine-to-serine mutant calpain baits (dCAPN, purple). (F) Coomassie-stained gel showing initial, his-tag purification of CAPN1[C115S]-3xFlag-6xHis (dCalpain-I) and CAPN2[C105S]-3xFlag-6xHis (dCalpain-II) in complex with CAPNS1(86-268) from *E. coli* (In- lysate input, FT- bead flow through, W- bead wash, Elu- elution from Ni^2+^ with 300 mM imidazole). Arrows indicate calpain proteins.

**S5.**
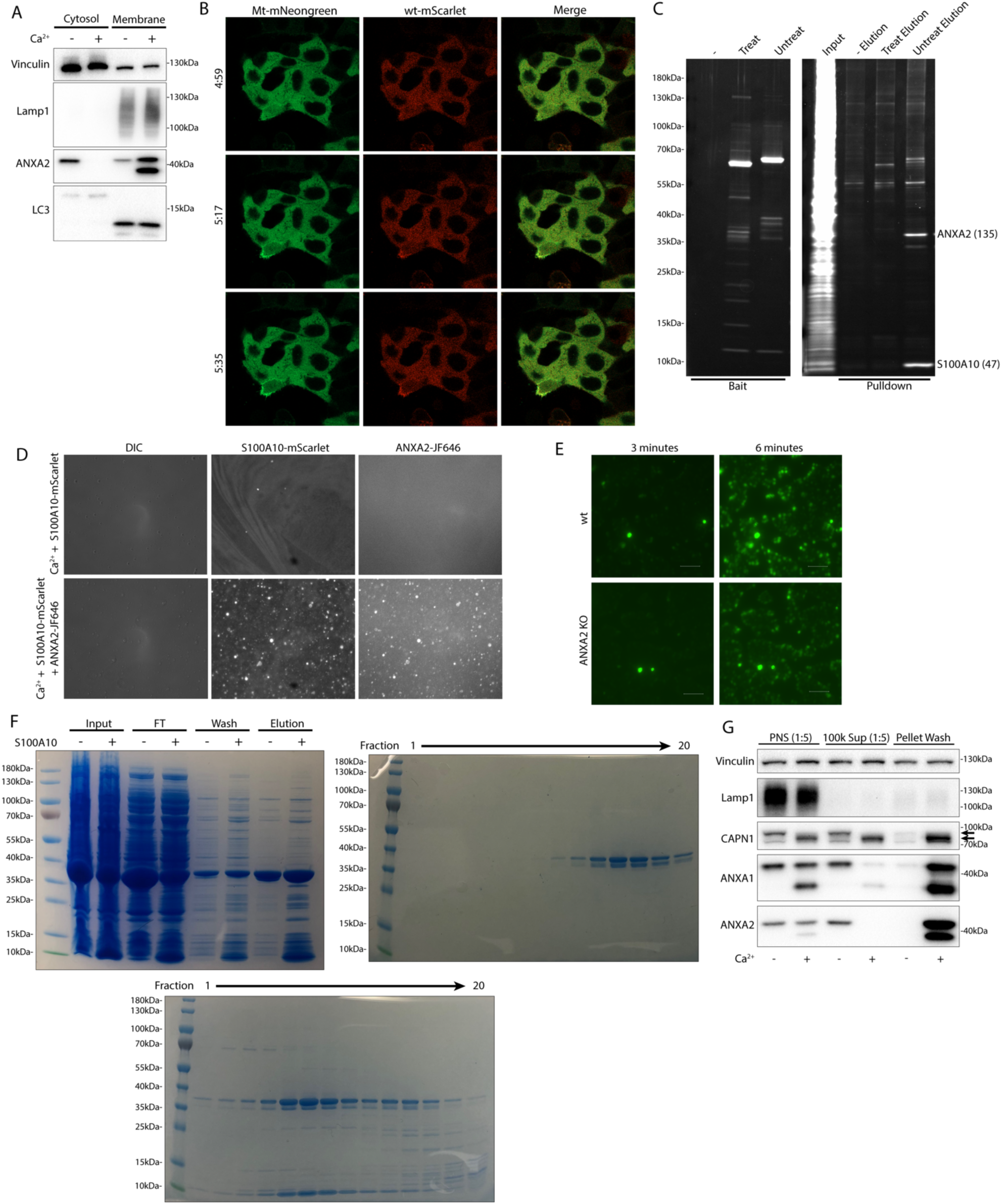
Calpain cleavage attenuates the membrane binding and scabbing activity of annexin A2. (A) Immunoblot analysis of cytosol and membrane factions, with or without 1 mM Ca^2+^ added prior to fractionation. (B) Representative confocal micrographs of ANXA2-mScarlet (wt-mScarlet) or ANXA2[P27D, P28D]-mNeonGreen (Mt-mScarlet)-expressing cells. Image times are relative to the addition of 400 ng/mL SLO. (C) Total protein (Sypro Ruby staining) analysis of HaloTag “pulldown” is shown, using no bait (−), porcine calpain-1-treated annexin A2-haloTag bait (Treat), or untreated annexin A2-haloTag bait (Untreat). Proteins are labeled with spectral counts, in parentheses, from gel excision-mass spectrometry. (D) Representative widefield micrographs of purified S100A10-mScarlet (1 μM) binding to GUVs with or without annexin A2-HaloTag-JF646 (1 μM). (E) Representative widefield micrographs of wildtype or annexin A2 KO cells treated with 200 ng/ml SLO and 2.5 μM Sytox Green without Ca^2+^ in the media. (F) Coomassie-stained gels showing purification of annexin A2 or annexin A2-S100A10 complex from *E. coli*. Panel I shows the initial purification by eluting annexin A2 from the *E. coli* lysate pellet fraction in a Ca^2+^-dependent manner (Input-lysate input, FT-100k x g pellet flow through, Wash-CaCl_2_-containing 100k x g pellet wash, Elution-10mM ATP elution from 100k x g pellet fraction). Where indicated lysate from S100A10-expressing *E. coli* was mixed 1:1 with annexin A2 *E. coli* lysate before purification. Panel II and III show size exclusion chromatography fractions of annexin A2 and annexin A2-S100A10 complex, respectively. (G) Immunoblot analysis shows mechanically lysed cells sequentially fractionated into post nuclear supernatant (PNS, diluted 1:5 before loading), 100k x g centrifuged PNS supernatant (100k Sup, diluted 1:5 before loading), and 100k x g pellet washed with 5 mM EGTA and recentrifuged (Pellet Wash). Samples where 1 mM Ca^2+^ was initially added to PNS are indicated. Arrows indicate uncleaved and autolyzed CAPN1.

